# Theory of Neuronal Perturbome: Linking Connectivity to Coding via Perturbations

**DOI:** 10.1101/2020.02.20.954222

**Authors:** Sadra Sadeh, Claudia Clopath

## Abstract

To unravel the functional properties of the brain, we need to untangle how neurons interact with each other and coordinate in large-scale recurrent networks. One way to address this question is to measure the functional influence of individual neurons on each other by perturbing them *in vivo*. Application of such single-neuron perturbations in mouse visual cortex has recently revealed feature-specific *suppression* between excitatory neurons, despite the presence of highly specific excitatory connectivity, which was deemed to underlie feature-specific *amplification*. Here, we studied which connectivity profiles are consistent with these seemingly contradictory observations, by modelling the effect of single-neuron perturbations in large-scale neuronal networks. Our numerical simulations and mathematical analysis revealed that, contrary to the prima facie assumption, neither inhibition-dominance nor broad inhibition alone were sufficient to explain the experimental findings; instead, strong and functionally specific excitatory-inhibitory connectivity was necessary, consistent with recent findings in the primary visual cortex of rodents. Such networks had a higher capacity to encode and decode natural images in turn, which was accompanied by the emergence of response gain nonlinearities at the population level. Our study provides a general computational framework to investigate how single-neuron perturbations are linked to cortical connectivity and sensory coding, and paves the road to map the *perturbome* of neuronal networks in future studies.

## Introduction

Perturbative approaches to study neuronal dynamics are becoming pivotal in our understanding of the brain’s function and dysfunction (Fenno, Yizhar and Deisseroth, 2011; Yizhar *et al.*, 2011; Boyden, 2015). They however often involve perturbation of a large number of neurons, which renders the analysis of the underlying circuitry challenging. A more simplified approach which has been pursued recently is to map the functional influence of individual neurons by perturbing a single neuron at a time. Such single-neuron perturbations have recently revealed feature-specific suppression between excitatory neurons in mouse visual cortex (Chettih and Harvey, 2019). But we still lack a mechanistic account of how these single-neuron functional influences are connected to cortical connectivity and dynamics, and how they can shed light on functional processing of realistic stimuli in large-scale cortical networks.

Specifically, how different motifs of excitatory (E) and inhibitory (I) connectivity interact with each other to give rise to functional properties of neuronal networks, and how this is manifested in single-neuron perturbations, remains unclear. For instance, several experimental studies have recently reported a highly specific pattern of connectivity in mouse primary visual cortex (V1), where excitatory neurons with similar functional properties (e.g. orientation-selectivity) are connected together with higher probability and with stronger weights (Ko *et al.*, 2011, 2013; Cossell *et al.*, 2015; Lee *et al.*, 2016). This was suggested to give rise to feature-specific *amplification* of the feedforward input by the recurrent network (Li *et al.*, 2013; Lien and Scanziani, 2013). The results of single-neuron perturbations, on the other hand, suggest that feature-specific suppression, rather than amplification, is the dominant mode of functional interaction between excitatory neurons (Chettih and Harvey, 2019). It has, therefore, remained puzzling how these seemingly paradoxical results should be interpreted and reconciled.

Here, we developed a theory of single-neuron perturbations and used computational modelling to shed light on these questions. We built and analyzed large-scale models of neuronal networks constrained with realistic receptive fields and experimentally reported motifs of recurrent connectivity and studied the effect of single-neuron perturbations in these networks. Specifically, we asked which cortical connectivity regimes are consistent with the experimental results of single-neuron perturbations. Our results highlighted the crucial role of inhibitory connectivity patterns, and how they interact with excitatory motifs to give rise to feature-specific effects (e.g. amplification and suppression). We found that to obtain feature-specific suppression, strong and functionally-specific subnetworks of E and I were necessary. That is, both E and I neurons with similar receptive fields (RFs) should be connected together more strongly than their non-similar counterparts, which was consistent with recent results in visual cortex (Znamenskiy *et al.*, 2018).

Our modelling results shed light on the above mentioned controversy by showing that feature-specific amplification *and* suppression could both exist in the cortex, depending on the regime of functional similarity between the influencers and the influencees. Our model suggests specific predictions on how to observe this in the cortex. Computational modelling also helped us to formulate further predictions that experiments could not directly assess, for instance regarding the temporal evolution of functional influence. We linked the result of single-neuron perturbations to sensory processing, by studying how our model networks in different regimes encode and decode natural images. More generally, we show that our theory can be extended to study multiple-cell perturbations to map the *perturbome* of neuronal networks in future.

## Results

### Single-Neuron Perturbations in Large-scale Networks of Visual Cortex

We studied the effect of single-neuron perturbations on functional properties of neurons in large-scale network models of visual cortex (**Figure 1A**). Individual excitatory and inhibitory neurons were modelled by two-dimensional visual receptive fields (RF) with randomly assigned initial parameters (e.g. preferred orientations and spatial frequencies) (**Figure 1B**; **Methods**). In accordance with experimental findings (Cossell *et al.*, 2015), the connectivity of neurons in the network was governed by a RF-similarity based rule, where neurons with more similar RFs had stronger connection weights (**Figures 1C,D**). We first simulated the responses of the network in the baseline state (i.e., before perturbation) (**Figure 1A**, upper), in response to gratings of different orientations and spatial frequencies (**Figure 1E**). We then simulated the response of the network with an extra perturbation of a single excitatory neuron (“influencer”) (**Figure 1A**, lower) and measured the change in the activity of other excitatory neurons in the network (“influencees”) (**Figure 1F**). The average response change of each influencee as a result of perturbation normalized by the strength of perturbation was taken as a measure of the functional “influence” of the single-neuron perturbation (**Figure 1G** and **Figure S1**).

**Figure 1.**
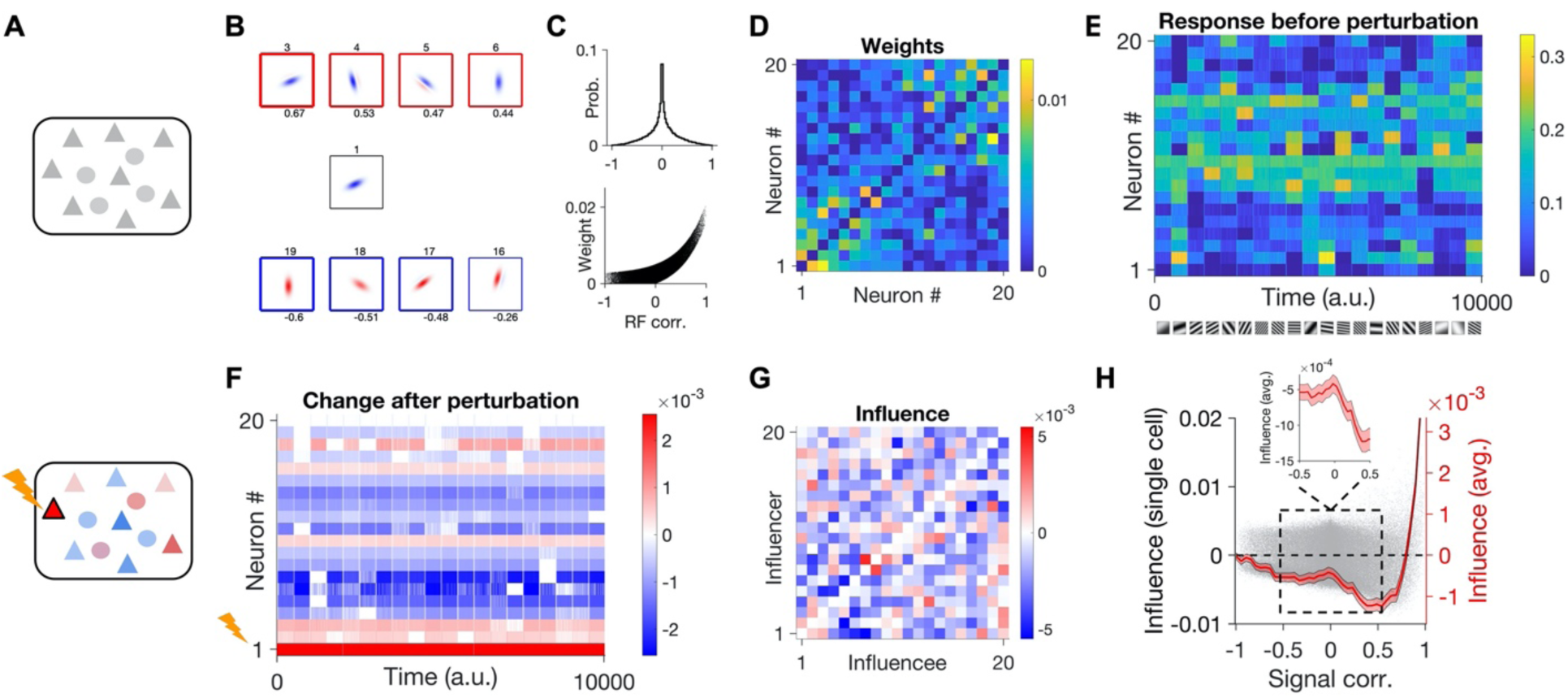
Influence of Single-Neuron Perturbations in Large-Scale Neuronal Networks. (A) Large-scale networks composed of excitatory (triangles) or inhibitory (circles) neurons are simulated in the baseline state (above) or after perturbing a single neuron (lower). (B) Example visual receptive fields (RFs) of excitatory neurons. Sample neurons with positive and negative RF correlations (CC) with the RF in the center are shown on the upper and lower rows (in red and blue squares), respectively. The thickness of lines around each RF is proportional to the absolute value of RF CC (indicated on the bottom). Examples RFs are 9 samples from 20 neurons (from the total of 400 excitatory units), which are shown throughout the figure. IDs of example neurons (indicated on top of each RF here) are the same in the rest of the figure, for comparison. (C) Distribution of RF correlations for all excitatory pairs in the network (upper), and the relationship between connection weights and RF correlations for the respective pairs (lower). (D) Sample weight matrix for 20 example excitatory neurons. Neurons are sorted according to their similarity (RF CC) to RF #1. (E) Firing rate response of example neurons to example static gratings (shown on the bottom). (F) Change in the response of neurons after perturbing neuron #1. (G) Influence (average response change normalized by the perturbation size) for all pairs of example excitatory neurons as influencers (different rows) or influencees (different columns). (H) Influence as a function of signal correlation for all excitatory pairs (gray dots) in the network. The average influence at different levels of signal correlation is plotted on the right (bin size: 0.05). Shading denotes ±sem. Inset: Zoom into the intermediate range of RF similarity.

To investigate how the interaction between neurons depends on their similarity, we plotted the influence for each pair of neurons (influencers and influencees) against their signal correlation (**Figure 1H**; see **Methods**). For moderate correlations, the net influence was negative, consistent with the average negative effect of single-neuron perturbations in experiments (Chettih and Harvey, 2019). Moreover, we observed “feature-specific suppression” in this regime, that is the negative influence was stronger for pairs with more similar response properties, on average (**Figure 1H**, inset). These results are consistent with feature-specific suppression observed in single-neuron perturbations *in vivo* (Chettih and Harvey, 2019).

However, this behavior changed for pairs with very strong response correlations, where we observed a positive influence, on average (**Figure 1H**). A similar trend had been observed for high “trace correlations” in the experiments (c.f. Fig. 5b in Chettih and Harvey, 2019). Based on our results, this regime of amplification is linked to RF similarity of neuronal pairs and hence can be assumed as “feature-specific amplification”. At the population level, these positive influences were stronger but less frequent, while the main bulk of influence between neuronal pairs was negative and small (**Figure S1**). These results therefore suggest different regimes of influence in the networks, whereby pairs of neurons with moderate response similarity (most pairs) show feature-specific *suppression* on average, while feature-specific *amplification* is dominant for highly similar RFs (rare examples).

### Cortical Connectivity and Single-Neuron Influence

To better understand how these feature-specific effects emerge, and how they are related to cortical connectivity, we developed a theoretical framework for the analysis of single-neuron perturbations in neuronal networks (**Figure 2**; see **Methods**). For linear networks, the theory can predict the impact of single-neuron perturbations on other neurons as a function of the weight matrix (**Methods**). We can, therefore, evaluate the average influence of neuronal pairs in the same networks as a function of their similarity. The theoretical prediction from the weight matrix shows the same non-monotonic behavior as our previous numerical simulations (**Figure 1H**), with feature-specific suppression for moderate response correlation and feature-specific amplification for highly similar RFs (**Figure 2A**). We therefore conclude that the main properties of feature-specific suppression/amplification arising from single-neuron perturbations can be deduced from neuronal interactions resulting from functional connectivity.

**Figure 2.**
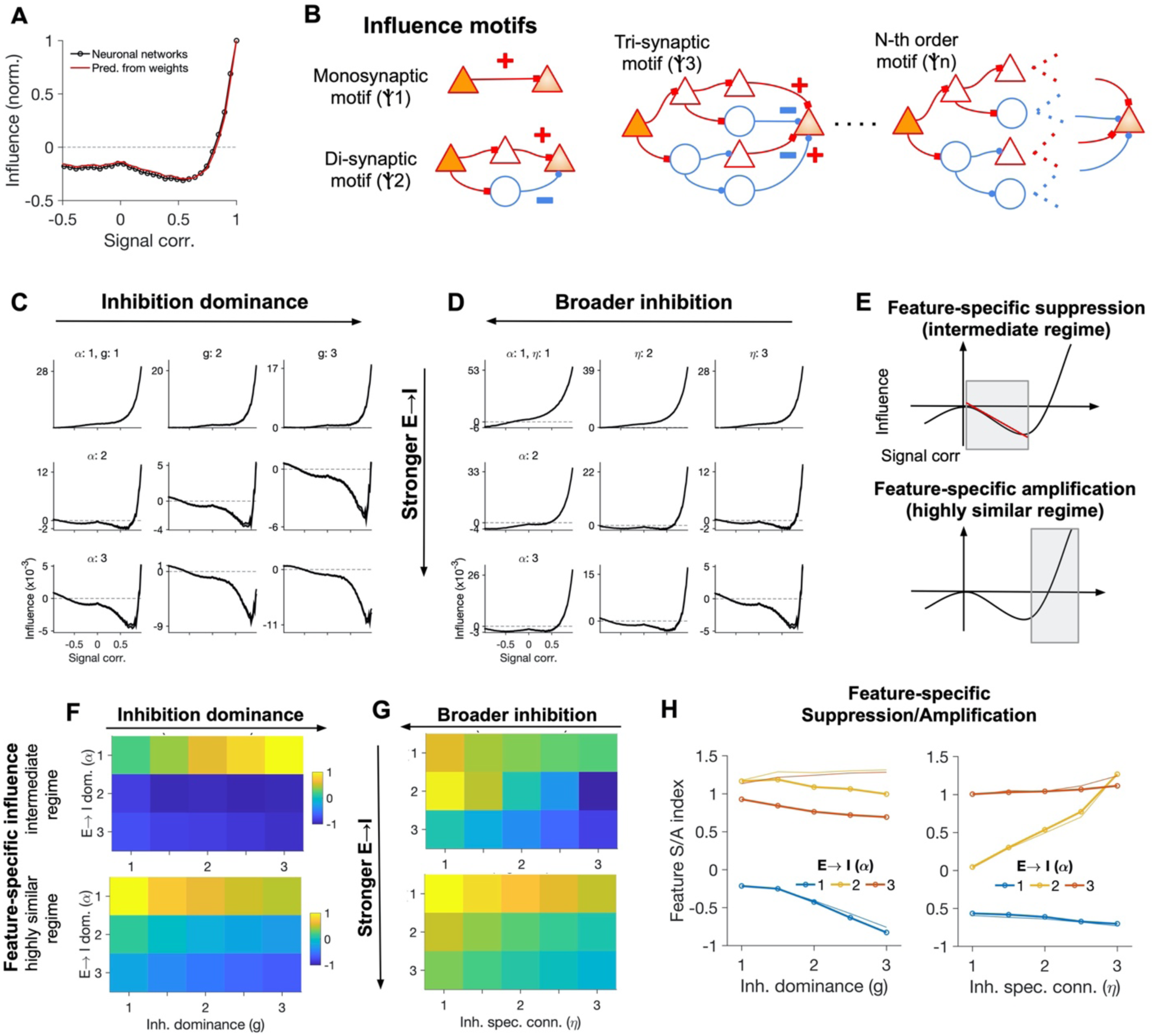
Connectivity Regimes and Feature-Specific Influence. (A) Average influence as a function of signal correlation in neuronal networks (as reported in **Figure 1H**) and the theoretical prediction of this influence from the weight matrix of the network. (B) Illustration of different motifs of influence (of different orders) mediating the influence of a perturbed excitatory neuron (influencer) on a postsynaptic excitatory target (influencee). The sign of the net influence at each branch is determined by considering the interaction of the signs of all synapses in the respective pathway. (C) The average influence as a functional of signal correlations inferred from the weight matrices of networks with different parameters of E→I connectivity and inhibition dominance. (D) Same as (C) for different strength of E→I connectivity and specificity of I→{E,I} connections. (E) Each pattern of influence was quantified by analyzing the degree of feature-specific amplification in the intermediate regime, quantified by the slope of the linear fit to the curve (upper) and the average amount of feature-specific amplification in the highly similar regime (lower). (F) Feature-specific influence for intermediate (upper; as described in **Figure 2E**, upper) and highly-similar regime (lower; as described in **Figure 2E**, lower) of neuronal similarity for different combination of inhibition-dominance (g) and strength of E→I connectivity in the network. (G) Same as (F) for different strength of E→I connectivity and specificity of I→{E,I} connections. (H) Feature-specific suppression/amplification (S/A) index, combining both aspects of suppression and amplification at different regimes, for different combination of parameters. Thicker lines show the values inferred from the weight matrix, and the thinner lines correspond to simulations of the neuronal networks (details of the results of rate-based simulations are shown in **Figure S2**).

While our analysis so far highlighted the key role of connectivity in feature-specific interactions, it does not reveal how different components of connectivity would give rise to suppressive or amplifying interactions. To shed light on this, we extended our mathematical framework to calculate the functional influence in terms of different pathways through which neurons can interact (“influence motifs”) (c.f. Pernice *et al.*, 2011) (**Figure 2B**; see **Methods**). The monosynaptic motif describes the direct pathway through which an excitatory neuron can influence another excitatory neuron. While the monosynaptic motif can only be excitatory, the di-synaptic motif confers positive as well as negative net influences (**Figure 2B**). Inhibition dominance (namely stronger inhibitory connections compared to E→E) can make the overall effect suppressive. However, too strong inhibition-dominance leads to a net positive influence as a result of disinhibition in the tri-synaptic motif, which can potentially counteract its suppressive effect in the di-synaptic motif. In strongly connected excitatory-inhibitory networks, the net effect of single-neuron perturbations can only be evaluated by accounting for all such higher-order motifs (**Figure 2B**).

We analytically calculated the net influence between two excitatory neurons by considering all possible excitatory and inhibitory motifs in between (**Methods**). The theory allowed us to evaluate the *prima facie* intuition that increasing the strength of inhibition alone is enough to mediate suppressive effects between excitatory neurons. Counterintuitively, our analysis revealed that inhibition-dominance alone cannot confer a net suppressive influence. Instead, increasing the weight of all inhibitory connections scales the positive influence of E neurons in a divisive manner (hence leading to a *divisive inhibition*; see **Methods**, Section 3.1.2). The direct weight of influence between two E neurons (J) would be scaled according to J/k (see **Methods, Eq. 35**), where the divisive term k increases with the overall strength of connectivity in the network, but also depends on inhibition-dominance (g). For g < 1, namely when inhibition is weaker than EE connections on average, k < 1, meaning that recurrent interactions in fact amplify the influence between a pair of EE neurons. For g > 1, that is when I connections are stronger than the EE weights, k > 1, hence the divisive scaling. The more the inhibition dominance (g), the larger the divisive term (k), and hence the smaller the influence of excitatory neurons on each other. However, the overall influence remains positive, which suggest that no net “negative” influence can result from single-neuron perturbations.

Our numerical simulation of functional influence in fact corroborated the above results: increasing the relative strength of inhibitory connections (parameterized by g) in networks with feature-specific connectivity did not introduce any feature-specific suppression (**Figure 2C**). However, when this was combined with strong E→I connections (parameterized by α), feature-specific suppression emerged. Feature-specific amplification for highly similar RFs was present for moderate values of g and α, but became less prominent for large values (**Figure 2C**). Our analytical results also suggested that broad inhibition alone does not confer feature-specific suppression (**Methods**), a finding which was further confirmed in simulations with broader inhibitory connectivity (parameterized by *η*, controlling the specificity of inhibitory connections) (**Figure 2D**). Simulation of neuronal networks with similar connection weights led to similar results, for both inhibition-dominance and broad inhibition scenarios (**Figure S2**).

To systematically characterize the functional properties of our networks, we developed an index which quantifies the simultaneous presence of feature-specific suppressions at intermediate regimes and feature-specific amplification for highly similar regimes (**Figure 2E**), as we described before (Feature S/A index; see **Methods**). Higher values of this index correspond to stronger feature-specific suppression at moderate levels of RF similarity *and* feature-specific amplification for highly similar RFs (**Figures 2F,G**). Consistent with our qualitative observation before (**Figures 2C,D**), neither inhibition-dominance nor broad connectivity of inhibition did result in high values of this index, in the absence of strong E→I connectivity (**Figure 2H**). Quantifying the functional behavior of rate-based networks with different parameters (**Figure S2**) led to very similar results (**Figure 2H**). Strong and specific E→I condition for the emergence of feature-specific suppression was in fact inferred from our analytical calculations (**Methods**), where *α* > 1 (i.e. stronger E→I connections compared to E→E weights) was necessary for a negative influence (**Eq. 80** in **Methods**). Our analysis thus captures the main functional behavior of neuronal networks as a result of single-neuron perturbations. These results suggest that strong and specific inhibition and E→I connectivity are necessary and sufficient conditions to obtain patterns of functional influence similar to the experimental results.

### Influence as a Function of Individual Features of Receptive Fields

We presented the results of single-neuron perturbations in terms of similarity of neuronal responses, as we had access to actual RFs of neurons in our model networks. However, mapping the full RF of neurons in experiments is not always feasible, and experimental results are often expressed in terms of marginal feature selectivity of neurons (e.g. their tuning to individual features of RFs like preferred orientation or spatial frequency). To relate better our results to such experiments (e.g. as in Chettih and Harvey, 2019), we analyzed the influence as a function of individual features of neurons (**Figure 3**).

**Figure 3.**
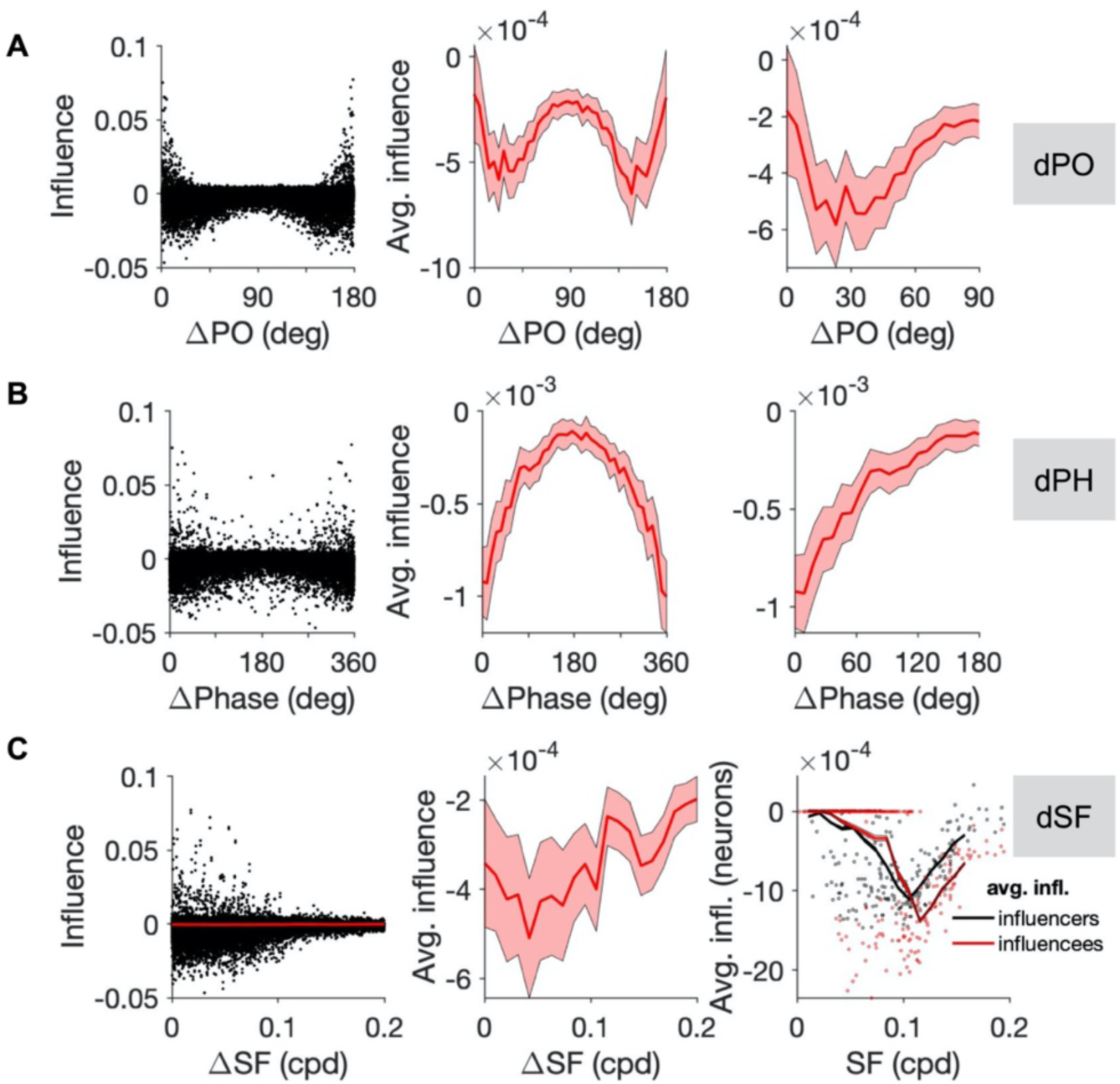
Influence as a Function of Individual Features. (A) Influence (inferred from perturbing neuronal networks similar to **Figure 1**) as a function of the difference in the preferred orientation (dPO) of neuronal pairs. Left: Distribution of all pairs. Middle: Average influence as a function of dPO (in bins of 4.5 degrees). Right: Zoom in (x-axis) of the average influence to highlight the nonmonotonic pattern. (B) Same as (A) for the difference in preferred spatial phase (dPhase). Bin size: 9°. (C) Influence as a function of the difference in the preferred spatial frequency (dSF) of neuronal pairs. Left: Distribution of all pairs (black) and the average influence (red). Middle: Average influence as a function of dSF (in bins of 4.5°). Right: Average influence per neuron (resulting from perturbing the neuron as an influencer (black) or observed by the neuron as an influencee (red)), as a function of its preferred SF. Dots show the result for all neurons in the network, and lines denote the average at each SF. Bin width: 0.01.

#### Preferred Orientation

Characterization of the influence as a function of the difference between the preferred orientation (PO) of the influencers and influencees revealed feature-specific suppression for intermediate PO differences (dPOs), where more suppression was observed between pairs with more similar POs (**Figure 3A**), in keeping with experimental results (c.f. Fig. 3i Chettih and Harvey, 2019). However, our analysis revealed an opposite trend of feature-specific amplification (i.e. less suppression for pairs with more similar POs) for very small differences (**Figure 3A**). That is a consequence of feature-specific amplification for the regime of highly similar RFs as we described before (**Figure 1H** and **Figure 2**), when that similarity is projected over an individual feature of RFs, namely their PO. Our results thus suggest that mapping the dPO of neuronal pairs with more resolution and/or larger sample size should reveal another regime of amplification, in addition to feature-specific suppression for the intermediate range.

#### Spatial Phase

Among the individual features of the RFs, we found that the preferred spatial phase of neuronal pairs revealed the strongest feature-specific suppression. Plotting the influence as a function of the difference in preferred spatial phase (dPH) of neuronal pairs revealed a monotonic increase with dPH, indicating that neuronal pairs with the closest preferred spatial phase show the most feature-specific suppression, on average (**Figure 3B**). Based on these results, we predict that analysis of influence as a function of phase difference could be the most significant predictor of the influence among the individual features of neuronal tuning.

#### Spatial Frequency

We also analyzed the dependence of influence on the difference in the preferred spatial frequency (dSF) of neuronal pairs (**Figure 3C**). Here, the relationship was less obvious and more noisy, especially for small to moderate dSFs (<0.1, **Figure 3C**). This is consistent with the experimental results, which did not reveal a significant dependence of feature-specific suppression on dSF. However, we found a significant bandpass dependence of influence, when we calculated the *average* influence for each neuron, either as an influencer (i.e., the average influence resulting from the neuron to all influencees) or as an influencee (i.e., the average influence experienced by the neuron from all influencers), as a function of the neuron’s preferred SF (**Figure 3C**).

#### Interaction of Individual Features

We next analyzed how the *interaction* of above mentioned features is related to the influence. That is, instead of analyzing the influence as a function of a single feature, we studied its changes in the space of multiple features (**Figure S3**). We analyzed the influence as a function of the conjoint distribution of differences in PO and phase (**Figure S3A**) or spatial frequency (**Figure S3B**). The bandpass dependence of influence on dPO that we observed before (**Figure 3A**) was exacerbated for small differences in both cases (small dPH and dSF regimes), and vanished for the regimes with large differences (**Figures S3A,B**). This suggests that controlling for the difference in other properties of the cells (individual features of their RFs here, e.g. SF or spatial phase), can amplify the effect of feature-specific influence for unique individual features. Analysis of the dependence of the influence on dPO, when controlling for dSF and dPH at the same time, revealed similar results (**Figure S3C**).

### Influence Resulting from Multiple-Neuron Perturbations

Our results in the previous section revealed how interaction of multiple features can shed light on additional properties of functional influence which could be masked when looking at single features individually. In this section, we asked how such interaction can be studied if, instead of single-neuron perturbations, the interactome of the network is mapped by multiple-neuron perturbations. To this end, we investigated the effect of double-neuron perturbations, in which two neurons are perturbed simultaneously to assay their combined influence on postsynaptic targets (**Figure 4A**). We repeated similar experiments as outlined before (e.g. in **Figure 1**) for such double-cell perturbations, and analyzed the influence as a function of the similarity of each influencee to both influencers (**Figure 4B**). Feature-specific suppression was evident for the moderate regime of RF similarity, especially for the region with moderate RF similarity to both influencers. Regions with the least RF similarity to one influencer, in contrast, revealed the most amplification when the RF similarity was high with regard to the other influencer (**Figure 4B**). Projecting the influence over a single dimension composed of both influencers revealed a stronger feature-specific suppression profile, when assayed as a function of RF similarity or an individual feature (preferred spatial phase) (**Figure 4C**).

**Figure 4:**
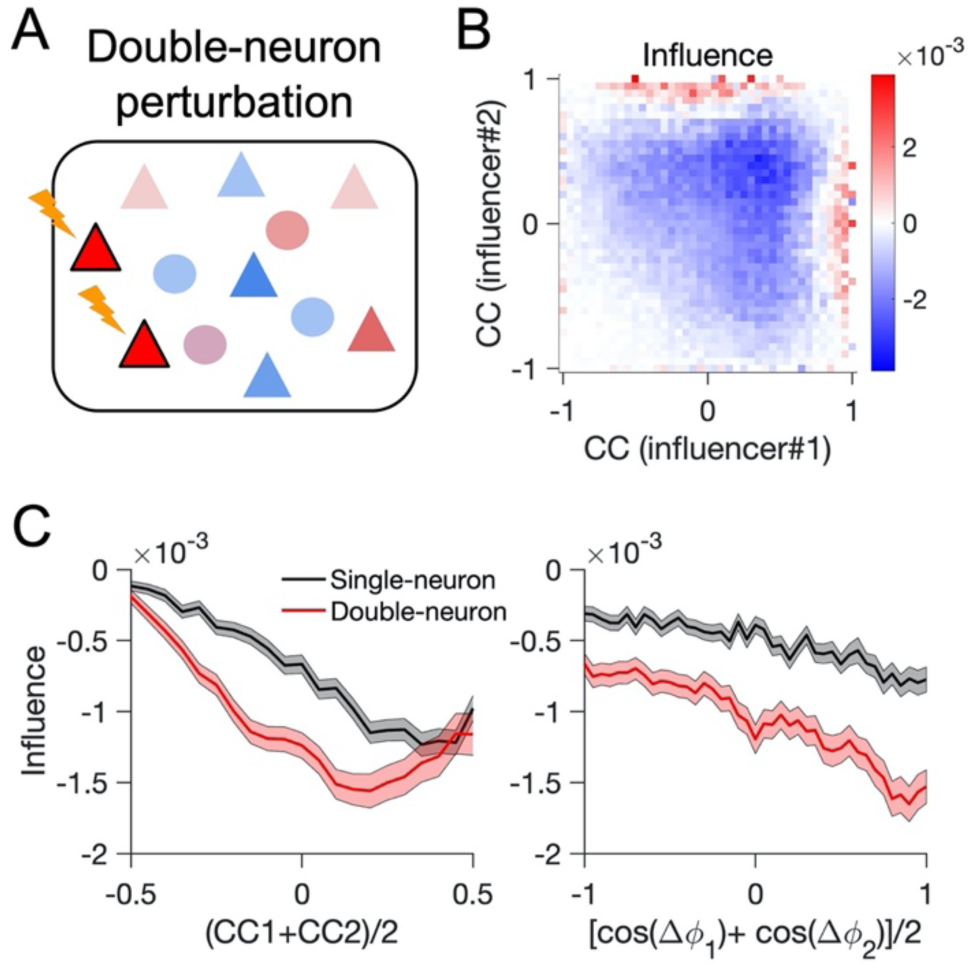
Mapping the Interaction of Influence in Double-Cell Perturbations. (A) Double-cell influence is assayed in neuronal networks by perturbing two excitatory neurons and quantifying the result of this dual perturbations on other neurons in the network. (B) Average influence as a function of the average response correlation of the two influencers with all the influencees. (C) Left: Average influence as a function of the average response correlation of the two influencers with influencees in double-neuron perturbations (red). CC1=CC2 for single-neuron perturbations (black). Right: Average influence as a function of the average difference in the spatial phases of the influencers with the influencees. Cosine of dPHs is used to obtain a normalized measure between −1 (most dissimilar) to 1 (most similar).

The interaction of influencers thus confers more feature-specific suppression on average. This interaction can be more systematically studied, by analyzing if the conjoint influence of two influencers is synergistic or antagonistic, namely whether the perturbation of neuron B in addition to neuron A increases or decreases their influence in isolation (**Figure S4A**). To analyze this we developed a “synergy index”, which quantifies if the change in double-neuron influences is amplifying or suppressing the single-neuron effects (see **Methods**). The synergy index is computed for each A-B-C triplet, where A is the first influencer neuron, B is the second neuron that is additionally perturbed, and C is the target influencee of single- and double-neuron perturbations. The average synergy over all target influencees (Cs) for a sample influencer A and all other second influencers (Bs) is shown in **Figure S4B**, as a function of the response correlation of A and Bs. The average synergy reveals a net positive synergy for all A-B pairs, but this effect is more prominent for A-B pairs with high response correlations. Similar trends were observed when we calculated such average synergy curves for other example influencers (As) in the network (**Figure S4C**). These results suggest that double-neuron (and, more generally, multiple-neuron) perturbations can be employed in future experiments to map the perturbome of neuronal networks, by analyzing the synergy of their interactions.

### Temporal Evolution of Influence

In our previous results, we discarded the transient activity and evaluated the influence from neuronal responses in the stationary state. But transient responses can reveal important insights about the operation of neuronal networks, especially how neuronal interactions evolve over time to shape the influence. We therefore analyzed the influence as inferred from the average activity at different time intervals after single-cell perturbations (**Figure 5**). Analyzing the pattern of average influence revealed that feature-specific amplification for very high response correlations was evident from very early responses, which is consistent with their excitatory monosynaptic nature (**Figure 2B**). However, feature-specific suppression emerged and strengthened over time, arguing for the polysynaptic nature of this component of influence (**Figure 5A**). We did not observe such dynamics in networks with weak E→I connectivity (**Figure 5B**). Our numerical results thus shed light on the evolution of the influence over time and is consistent with our analytical results on the significance of higher-order motifs of interaction in the emergence of feature-specific suppression.

### Inhibitory Single-Neuron Influence

We also studied how single-neuron perturbations would change, if inhibitory neurons are perturbed as influencers instead of excitatory neurons. Mapping such inhibitory influences is more challenging experimentally, but we could investigate it in our model (**Figure S5**). We observed stronger negative influences, on average (**Figure S5**), presumably due to stronger weights of I→E connections. Beyond the mean suppression, a negative slope of influence versus response correlation (which indicates feature-specific suppression) was observed for higher signal correlations (**Figure S5A**), as opposed to excitatory single-cell perturbations where such a feature-specific suppression was present in the intermediate regime (illustrated in **Figure 2E**). In fact, lack of a significant negative slope for intermediate positive signal correlations indicated an absence of feature-specific suppression in this regime (**Figure S5B**). The negative slope was, however, present for negative signal correlations in this regime, leading to some degree of apparent feature-specific “*amplification”* for intermediate negative correlations (**Figure S5B**). Thus, the pattern of inhibitory influence in the intermediate regime seems to be the opposite of the pattern of excitatory influence (c.f. **Figure 1H**), while the feature-specific amplification for highly similar regime is obviously missing. Another conspicuous difference between excitatory and inhibitory single-neuron perturbations was in the transient responses of the latter, which did not show a significant change in overall shape over time (**Figure S5C**). This argues that polysynaptic motifs are more important in establishing feature-specific suppression in excitatory single-neuron perturbations, as it involves the interaction of excitation and inhibition.

### Functional Consequences for Sensory Processing

Our work so far revealed how different connectivity profiles lead to various patterns of feature-specific suppression/amplification when individual neurons are perturbed. But single neurons are rarely perturbed in isolation. Instead, the realistic operating regime of the brain involves collective activation of neurons, for instance in response to external stimuli. To examine the functional consequences of such single-cell properties in naturalistic conditions, we therefore need to study the response of populations of neurons in different regimes. To address this, we presented natural images to large-scale visual networks. The feedforward input was obtained by filtering the images by the RF of individual neurons, and the output of the network was read from the population activity (**Figure 6A**; see **Methods**). We then analyzed how different networks transformed the input to output in different regimes.

**Figure 5:**
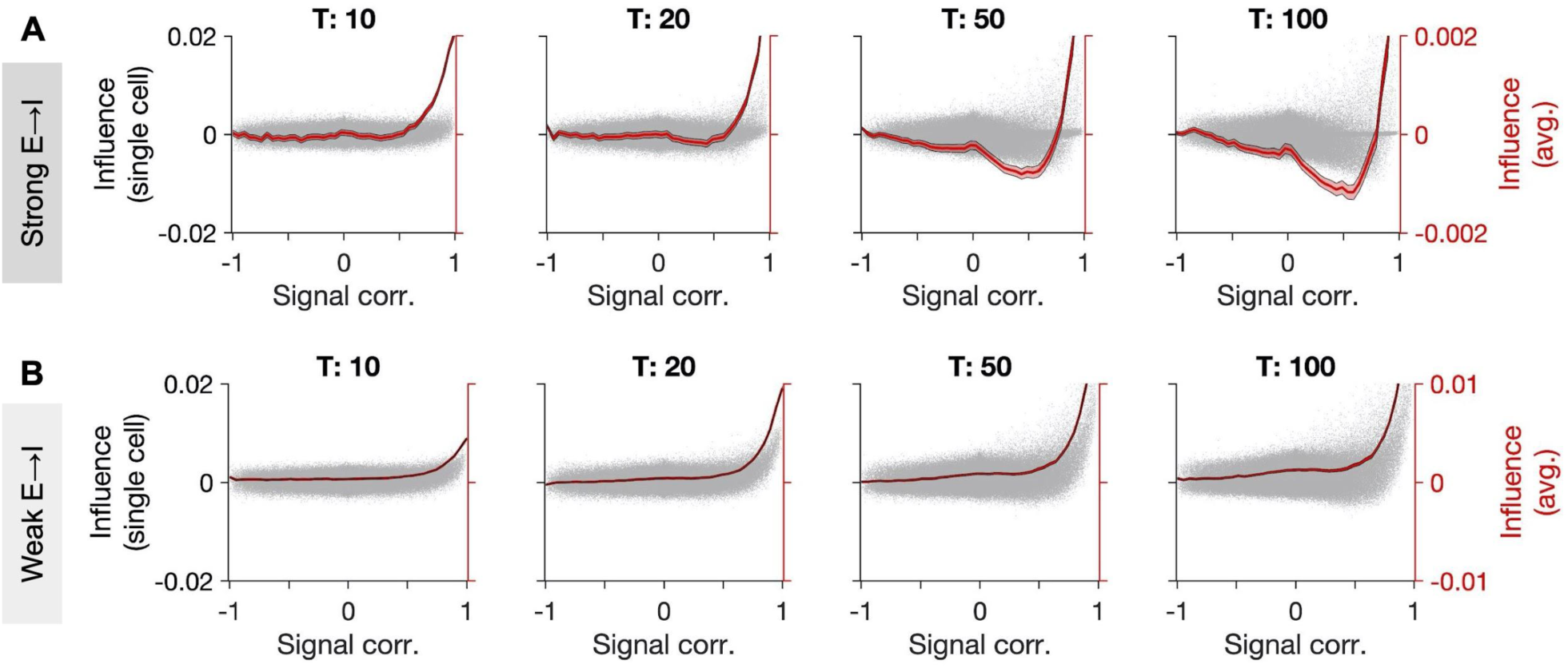
Temporal Evolution of Influence Inferred from Transient Responses. (A) Influence as a function of signal correlation in neuronal networks (similar to **Figure 1H**), when the influence is inferred from the average firing rate up to time T. Gray dots: all pairs; red: average influence (bin size: 0.05). Shading denotes ±sem at each bin. *α* = 2, *g* = 2, *η* = 3. (B) Same for a network with weaker E→I connectivity (*α* = 1).

**Figure 6.**
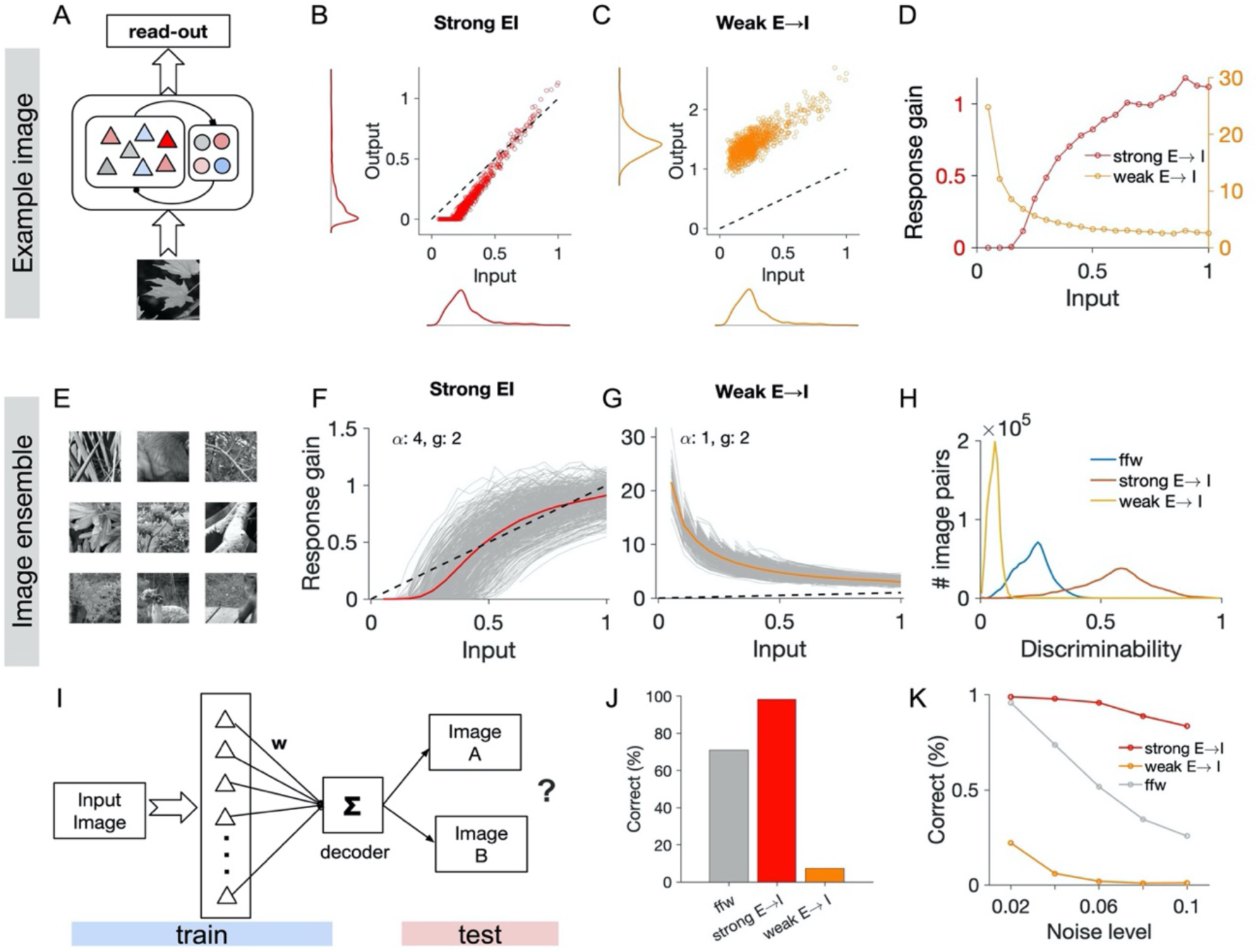
Population Responses in Different Regimes and Consequences for Sensory Processing. (A) The response of the network is obtained by projecting the input image over RF of neurons (feedforward input) and accounting for recurrent interactions resulting from the weight matrix. (B) Input and output of excitatory neurons in the network for a weight matrix with strong E→I. Input and output are both normalized to the maximum level of input in response to the image. Distribution of input and output are shown on respective axes. (C) Same as (B) for a weight matrix with weak E→I. (D) Response gain (output divided by input) for each excitatory neuron as a function of input, for weight matrices with weak and strong E→I, respectively. (E) Sample images from an ensemble of 617 natural images used to test the networks. (F) Individual response gain curves (as In (D)) for individual images (gray) and their average (red). (G) Same as in (F) for a weight matrix with weak E→I connections. (H) Discriminability of population responses, calculated as the normalized angle (by 90°) of the vectors of population responses to all pairs of natural images (see **Methods**). (I) For each image, a decoder is trained to distinguish the image from half of the images in the ensemble. It is then tested to distinguish the image from images in the other half of the ensemble. (J) Percentage of the correct responses of decoder in (I) in distinguishing the target image. (K) Percentage of correct responses at different levels of noise corrupting the population responses.

We first looked at networks similar to those with feature-specific suppression/amplification as a result of single-neuron perturbations (e.g. as in **Figure 1**). We analyzed the output activity of excitatory neurons as a function of their input in response to a sample image (**Figure 6B)**. For such networks, the activity of neurons with small inputs (small feedforward projections) was suppressed, while the activity of neurons with medium and large feedforward projections were mainly maintained or amplified (**Figure 6B**). To understand what underlies such a transformation, we repeated the input-output analysis for the same network but with weaker E-I connectivity. Here, we did not observe the same nonlinear transfer function; instead, output responses were generally amplified with respect to the input (**Figure 6C**).

To quantify the transformation, we calculated the response gain for each neuron as a factor with which the input needs to be multiplied to obtain the output. Neurons in the original networks (with strong E-I connections) showed a sigmoid response gain function: close to zero gains for small inputs and high gains for larger inputs, with a saturation trend at very high feedforward projections (**Figure 6D**). In contrast, the network with weak E-I connections showed the opposite trend: higher gains for neurons with small feedforward projections, and the least amplification for intermediate and large inputs (**Figure 6D**). We characterized such response gain curves for an ensemble of natural images (**Figure 6E**) and observed similar nonlinear behavior for both networks across the images (**Figures 6F,G**). These results suggest that recurrent interactions in the networks with strong E-E connections can amplify the “noise”, if the E-I interaction is weak; however, when strong and functionally specific connectivity exists between E-E *and* E-I connections, neuronal responses show a selective suppression of the noise and enhancement of the signal.

Lack of sigmoid-shape transfer functions in networks with weaker E-I connections can be a result of less inhibition in the network. We therefore asked if an unselective increase of inhibition in our networks can compensate for the weakening of E-I connections, and restore the transfer function. To test that, we studied networks with an increase in inhibition-dominance or broadness of the inhibitory connectivity, similar to the procedure we used before to evaluate different regimes of feature-specific suppression/amplification in these networks (c.f. **Figures 2C,D**). Under both scenarios, we observed qualitatively similar response gain curves as in networks with weak E-I connections, and sigmoid nonlinearity did not emerge as a result of nonspecific inhibition (**Figure S6**). These results corroborates that strong and functionally-specific E-I connections, which were necessary to obtain feature-specific suppression and amplification for different regimes, are also necessary for the emergence of sigmoid-shape nonlinear transfer functions, which can potentially enhance sensory coding.

To evaluate more directly the contribution of such nonlinear transfer functions to sensory processing, we studied the representation of natural images in different networks. We assessed how population responses to different images could be distinguished in networks with different nonlinearities. This was measured by quantifying the distance between population vectors in the N-dimensional space of neural activity (where N is the number of neurons). If the nonspecific component induced by all images over neurons is smaller than the specific projections, the distance between the vectors of population activity is small and hence it is more difficult to discriminate different images. On the other hand, orthogonal representations, namely patterns of activity with the least overlap, enable the most discriminability for image pairs. We quantified such discriminability of population representations for all pairs of natural images in different networks (**Methods**). The results revealed that networks with strong E-I connections increased the discriminability of feedforward projections, whereas networks with weak E-I had lower discriminability (**Figure 6H**). By suppressing the redundant information and enhancing the representation of more informative neurons, networks with the sigmoid nonlinearity can therefore provide a more efficient population code to represent natural images.

The enhancement in the encoding of visual stimuli should lead to better decoding capacities of visual networks. To test this directly, we assessed the capacity of different networks in distinguishing different natural images. We trained a decoder to discriminate a target image from other images in the ensemble, based on the population activity of excitatory neurons (**Methods**). We then tested the discrimination accuracy of the decoder when other non-target images (not seen during the training) were shown, under different levels of noise (**Figure 6I**; see **Methods**). Decoders which performed their discrimination based on the population activity of networks with strong E-I connections performed significantly better than decoders based on the feedforward input, while networks with weak E-I weights did much worse than both (**Figure 6J**). Increasing the level of noise reduced the accuracy for all networks, but networks with strong E-I connections outperformed other networks consistently; moreover, they showed the most robust behavior and were affected the least by noise (**Figure 6K**). We therefore conclude that nonlinear response gains, emerging in networks with strong E-I connections and feature-specific suppression/amplification in single-neuron perturbations, improve image processing by increasing the capacity of a downstream decoder to distinguish different stimuli.

## Discussion

We presented computational models and mathematical analysis of the functional effects of single-neuron perturbations in neuronal networks. Our results revealed specific connectivity motifs necessary for the emergence of feature-specific suppression and amplification for moderately-similar and highly-similar receptive fields. Strong and specific E-I connections were necessary in combination with strong inhibitory connectivity to yield those results. A key finding of our study is thus the importance of strong and specific E-I connectivity. This was necessary to explain the experimental results of single-neuron perturbations (Chettih and Harvey, 2019), but also consistent with recent experimental reports of the specificity of E→I and I→E connections in visual cortex (Znamenskiy *et al.*, 2018). Our theoretical analysis suggests that the most feature-specific suppression is achieved when these two motifs are balanced. Based on these results, an important implication of our study is that selective targeting of E-I connections, e.g. by optogenetics techniques, should modulate the feature-specific suppression.

We found that the same connectivity profiles also gave rise to the nonlinearity of neuronal responses, which underlie the enhancement of population capacity to discriminate natural images. A similar nonlinearity has been suggested to explain the visual responses to natural images in mouse V1 (Fig. 2g in Yoshida and Ohki, 2020). A prediction of our model is that such a sigmoid nonlinearity can be an emergent property of neuronal responses at the population level, as networks with different connectivity profiles expressed different nonlinearities in our simulations. Our model neurons in fact lacked such nonlinear transfer function at the single-cell level, and, given the operating regime of V1 (low firing rates and fluctuation-driven regimes of activity due to E-I balance), it seems unlikely that such a nonlinearity can arise from neuronal responses in isolation. Selective perturbation of the recurrent circuitry, especially targeting E-I connections, is thus needed in future studies to address the presence and nature of this nonlinearity, and its potential link to the representation capacity of the population code.

Strong E-I connectivity has been reported in many cortices across different species (Molnár *et al.*, 2008, 2016; Hofer *et al.*, 2011). Specifically, very large excitatory inputs from pyramidal cells to inhibitory neurons have been observed in humans, a property that has been absent in any other nonhuman cortices (Molnár *et al.*, 2008; Szegedi *et al.*, 2016), which may argue for the prominent role of this connectivity motif in complex cognitive processing. However, in the absence of specific mapping of their functional properties, and in view of the broad selectivity of inhibitory neurons (Bock *et al.*, 2011; Hofer *et al.*, 2011), it has been assumed that these connections are nonspecific and provide a blanket of inhibition to the local network (Fino and Yuste, 2011; Packer and Yuste, 2011; Karnani, Agetsuma and Yuste, 2014). Recent studies, on the other hand, have revealed the emergence of specific E-I subnetworks, emphasizing the potential significance of selective inhibition for cortical computation (Wilson *et al.*, 2017; Khan *et al.*, 2018; Najafi *et al.*, 2020). In mouse visual cortex, dendrite-targeting somatostatin positive (SOM+) inhibitory neurons have been reported to have comparable levels of orientation selectivity to excitatory neurons in both L4 and L23 (Ma *et al.*, 2010), and even within parvalbumin positive (PV+) interneurons a range of selectivity has been observed depending on the extent of their dendritic tree (Runyan and Sur, 2013). Future studies are thus needed to address the functional connectivity of E-I subnetworks more systematically to shed light on the specificity of E-I interactions. A case in point is a recent study of odor processing in the olfactory bulb of larval zebrafish (Wanner and Friedrich, 2020). By mapping the functional connectomics via dense reconstructions of wiring diagrams (as opposed to sparse sampling of connections), the study could shed light on the higher-order interactions of excitatory and inhibitory neurons. Interestingly, the study found that bidirectional E-I connectivity is implicated in the decorrelation of odor responses via feature suppression.

In addition to linking connectivity and coding in single-neuron perturbations, our theory outlines how multiple-cell perturbations can be used to study functional properties of neuronal networks, by mapping higher-order interactions between influencers. A similar mathematical approach has been recruited recently to analyze the interaction of drugs and their resulting changes in cell morphologies, in order to shed light on the link between drug combinations and treatment of diseases (Caldera *et al.*, 2019). Targeted multiple-neuron perturbations of functionally identified neurons have in fact been used recently to shed light on the dynamics of persistent activity and short-term memory in mice (Daie, Svoboda and Druckmann, 2019). Similar approaches can be recruited to reveal how neurons work in tandem to shape functional processing in sensory cortices, with the possibility that different network perturbomes can dissociate between functional and dysfunctional circuitries.

Our study also suggests that the temporal dynamics of the evolution of feature-specific suppression/amplification can reveal fundamental insights about the operation of the network. In our model neuronal networks, we found that feature-specific suppression emerges later than feature-specific amplification as a result of polysynaptic interactions (**Figure 5**). Testing if such a pattern also exists in the cortex would have implications for the connectivity and function of cortical networks (e.g. transient responses versus sustained activity). Although technically challenging, future experiments can use population voltage-based measurements (Knöpfel and Song, 2019; Piatkevich *et al.*, 2019) combined with single-neuron optogenetic perturbations to cast light on this important aspect. The temporal profile of functional influence can be further combined with multiple-neuron perturbations to map the temporal perturbome of neuronal networks, which will provide a more complete picture of functional and temporal patterns of processing in the brain.

In summary, our study provides a general mathematical framework to study the effect of single- (and multiple-) neuron perturbations in excitatory-inhibitory neuronal networks. By applying it to the visual cortex, we could unveil connectivity principles underlying the emergence of feature-specific influence in recent single-neuron perturbations, and predict further properties of visual networks. The model specifically provided an explanation for the mutual presence of functionally specific excitatory connectivity and feature-specific suppressive influence of perturbations, which seemed contradictory in view of previous experiments (Ko *et al.*, 2013; Cossell *et al.*, 2015; Lee *et al.*, 2016; Chettih and Harvey, 2019). The modelling framework can be used in future studies to link cortical connectivity and dynamics to function in perturbation experiments.

## Acknowledgement

We thank Arvind Kumar and Volker Pernice for comments on the manuscript. Code for reproducing main simulations and results will be available from ModelDB upon publication.

## Supplementary Figures

**Figure S1.**
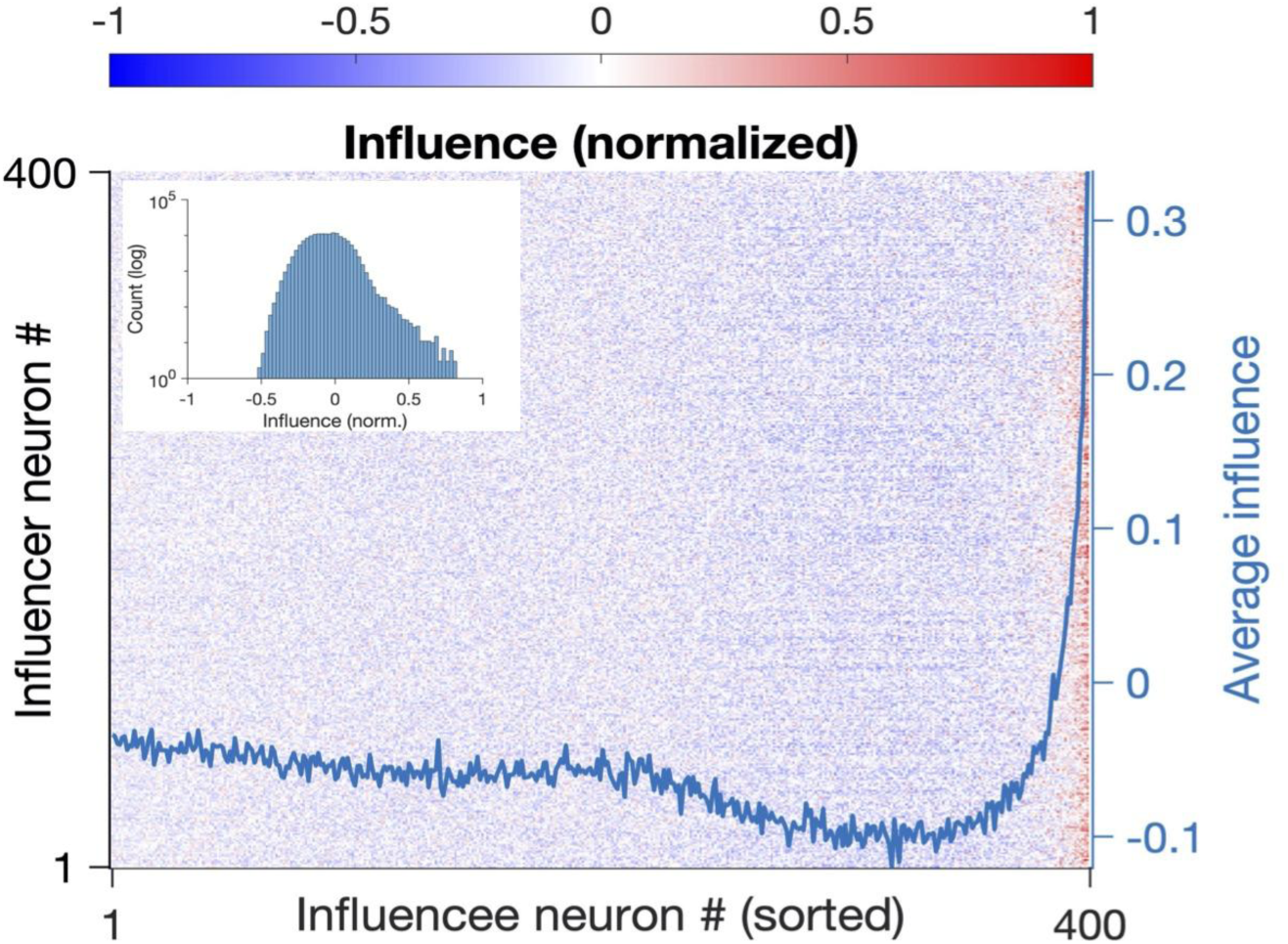
Influence for All Neurons in the Network (related to Figure 1). The matrix of normalized influence between influencers (different rows) and influencees (different columns) similar to **Figure 1G** but for all excitatory neurons in the network. Influence is normalized by the maximum absolute value of influence between all excitatory pairs. In each row, the influencees are sorted according to their response correlation with the respective influencer in an ascending order. The average influence for each column is plotted on the right. Inset: The distribution of normalized influence for all excitatory pairs in the network. Note the logarithmic scale on the y-axis.

**Figure S2:**
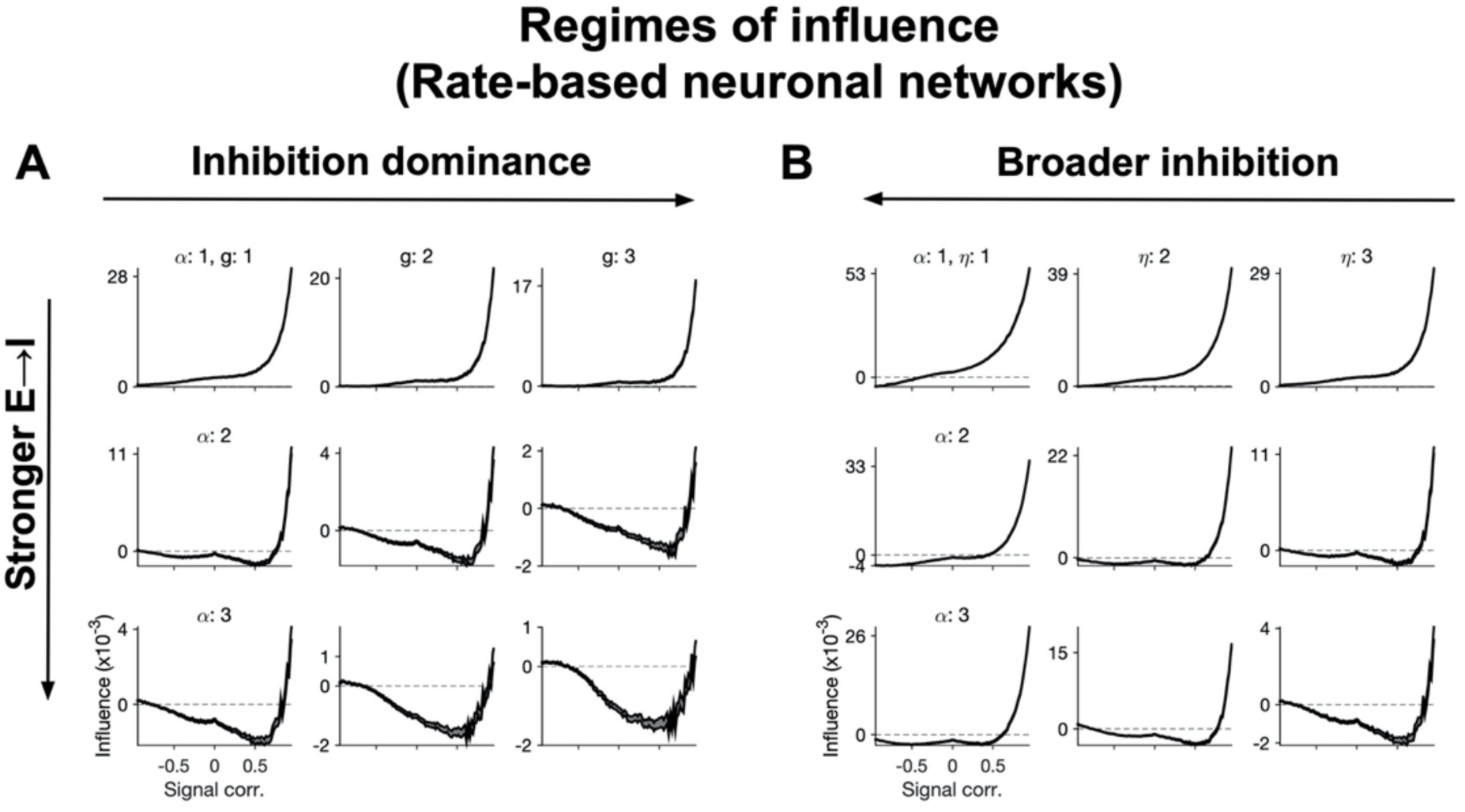
Regimes of Feature-Specific Influence in Neuronal Networks (related to Figure 2). (A,B) Same as **Figure 2C** and **Figure 2D**, respectively, for rate-based neuronal networks with similar weight matrices. For each network, the weight matrix is generated with the same combination of parameters as in **Figures 2C,D**, respectively, but instead of inferring the influence from the weight matrix (as in **Eq. 19**), the influence is calculated from rate-based simulations of the network (**Eq. 14**).

**Figure S3:**
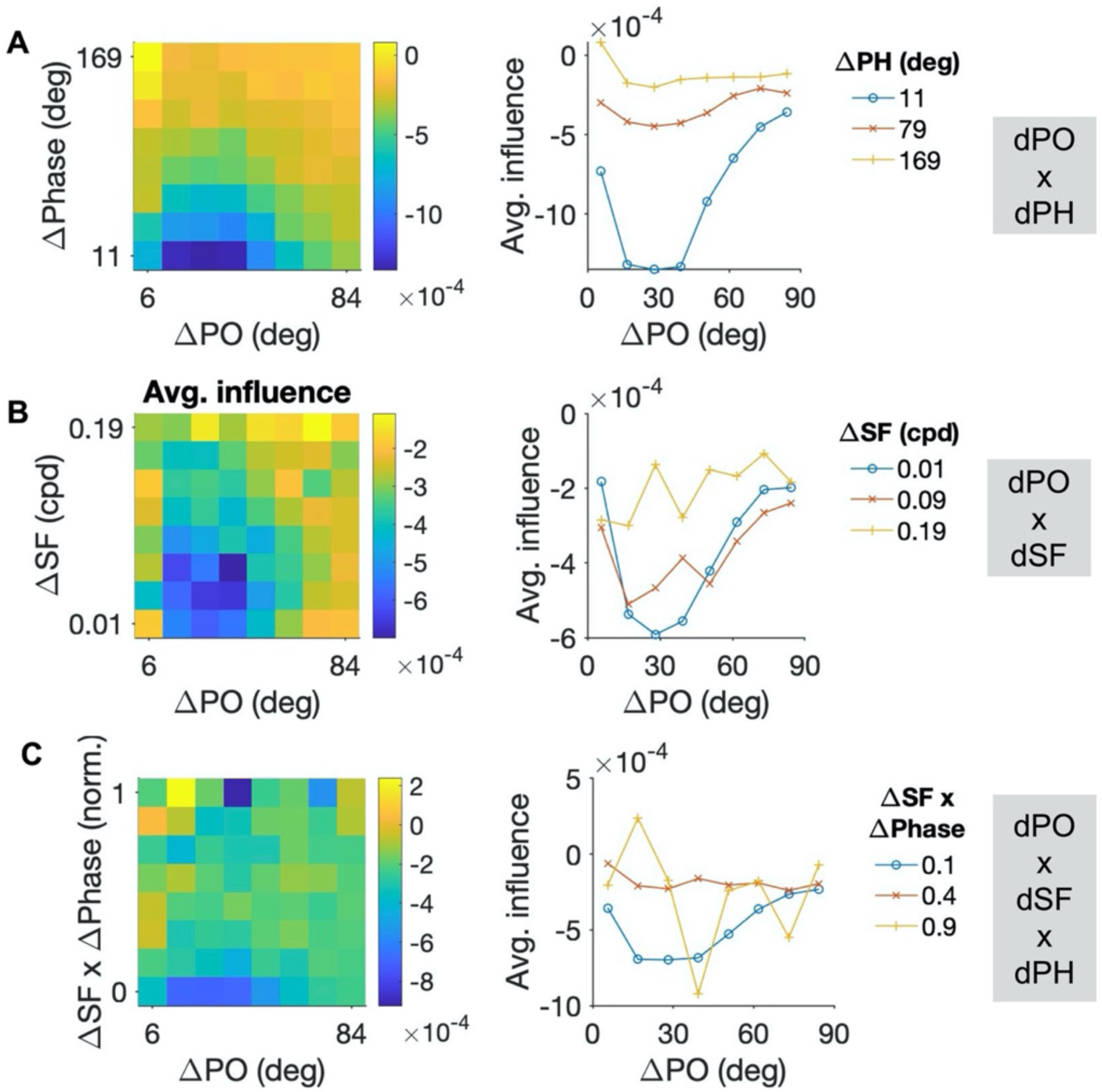
Influence as a Function of the Interaction of Individual Features (related to Figure 3). Same as **Figure 3**, but for interaction of individual features. (A) Left: Average influence for all pairs within a given range of dPO and dPH. Right: Average influence as a function of dPO for three levels (minimum, medium and maximum) of dPH. (B) Same as (D) for the interaction of dPO and dSF. (C) Same as (D,E) for the interaction of dPO with the conjoint change of SF and phase (dSF x dPhase). dSF and dPhase are both normalized to their maximum values, respectively, and then multiplied to obtain a single variable.

**Figure S4:**
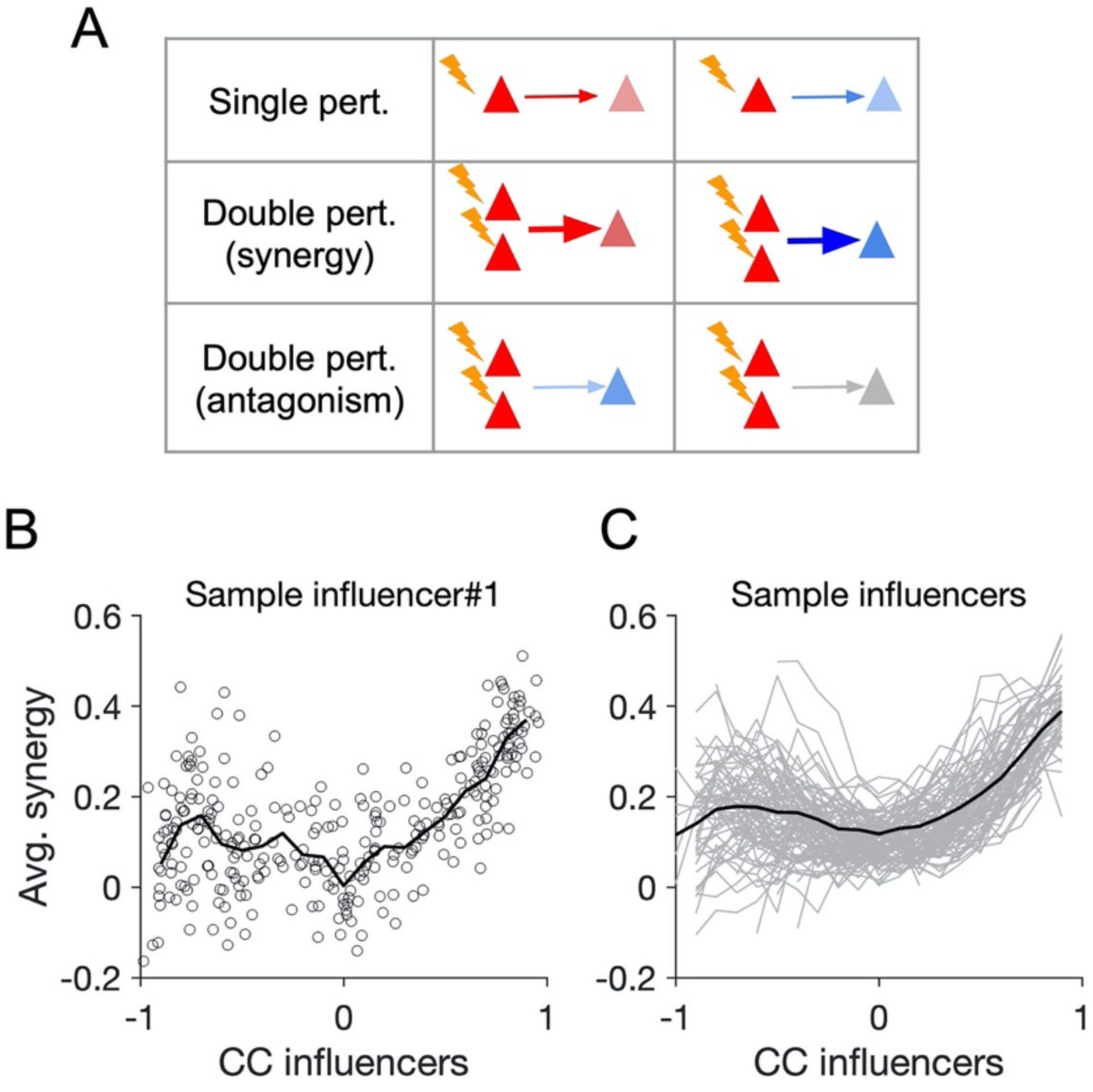
Synergy of Interaction in Double-Cell Perturbations (related to Figure 4). (A) Illustration of different outcomes of double-neuron perturbations compared to single-neuron results. First column: An influencer with a net positive influence on an influencee (first row) can experience synergistic interaction with another influencer if the net influence of the double-neuron perturbation is more positive (second row), or an antagonistic interaction if the net influence is less positive or negative (third row). Second column: Example of synergistic or antagonistic interaction for a negative single-neuron influence. (B) Average synergy index (see **Methods**) of an example first influencer with all other influencees as a function of response correlation of the first influencer with the second influencers. Black line shows the average in each CC bin (bin width: 0.1). (C) The average synergy as a function of response correlation (black line in (B)) for 100 sample first influencers (gray). Black line shows the average across all curves. Bin width: 0.1.

**Figure S5:**
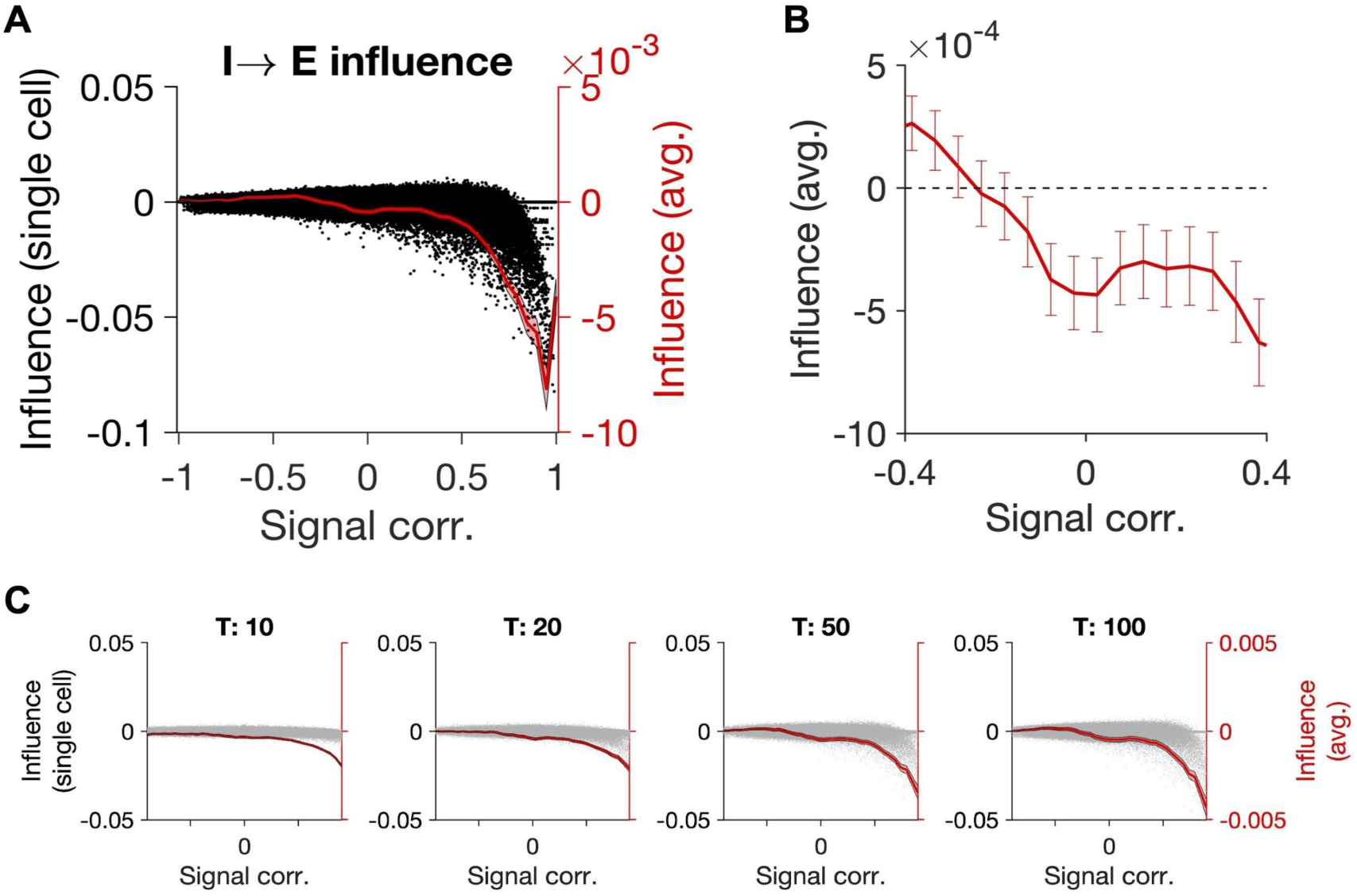
Influence Inferred from Perturbing Single Inhibitory Neurons (related to Figure 5). (A) Influence of inhibitory neurons on excitatory neurons for all pairs (black) and as average calculated for all pairs within a given range of signal correlation (bin size: 0.05). (B) Zoom in for the intermediate range. Error bars denote ±sem. (C) Same as in **Figure 5** for inhibitory single-neuron perturbations.

**Figure S6:**
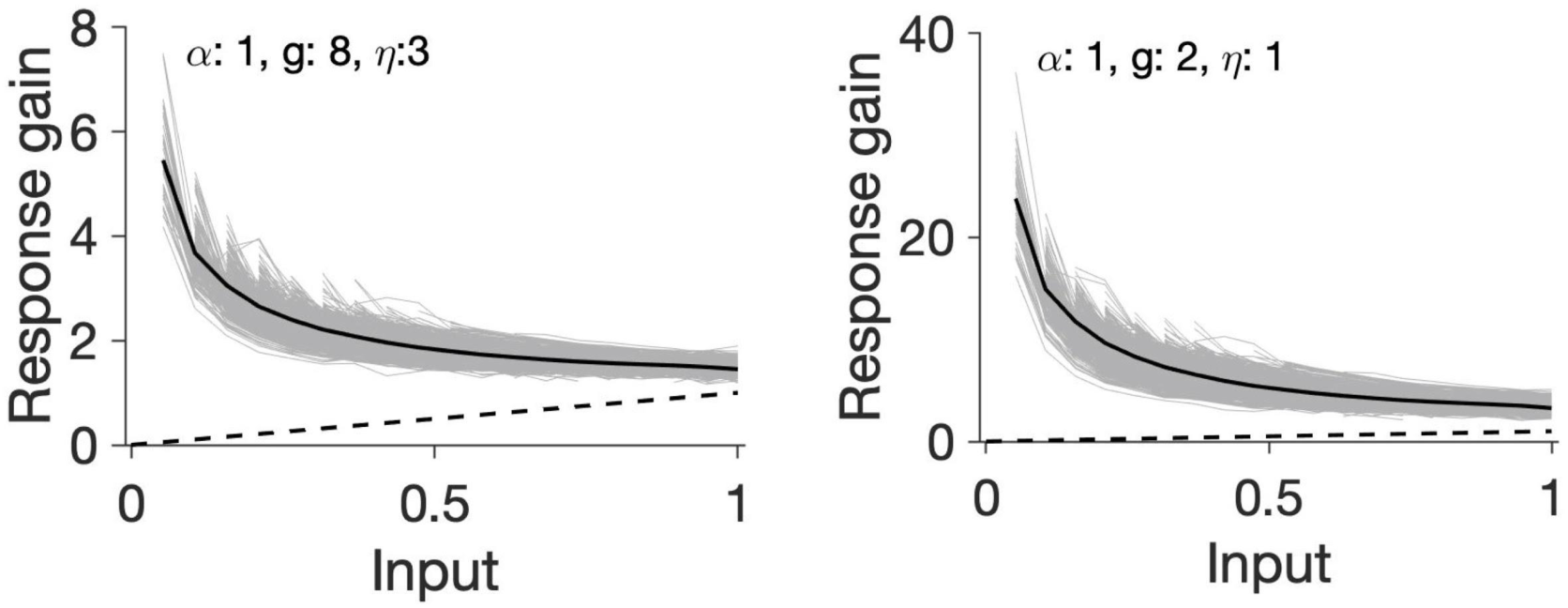
Response Gain Curves for Networks with Weak E→I and Stronger or Broader Inhibition (related to Figure 6). Same as **Figure 6G**, for control conditions where the network with weak E→I connections had either higher inhibition dominance (left) or broader inhibition (right). Neither of the two conditions resulted in sigmoid nonlinearity of gains as observed in **Figure 6F**.

## Methods

### 1 Network simulations

#### 1.1 Neuronal receptive fields

Visual receptive field (RF) of neurons were modeled as two-dimensional Gabor fields:

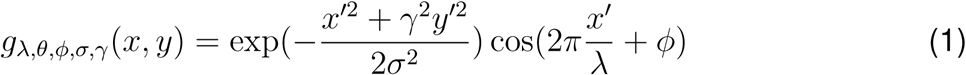

where

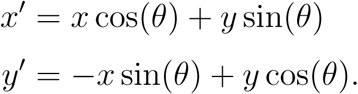

*x* and *y* are the position on the visual field, *σ* sets the size, *θ* is the preferred orientation, *ω* = 1*/*λ the spatial frequency, and *ϕ* the spatial phase of the receptive field. RFs are instantiated on a *L* × *L* field, with their resolutions expressed in pixels per degree (ppd), and centered at (*x*_0_, *y*_0_). Unless stated otherwise, parameters are chosen as the default values below: *L* = 50, ppd = 4, *σ* = 2.5, *γ* = 0.5. The following parameters are drawn randomly from a uniform distribution: (*x*_0_, *y*_0_) from [−1.25, 1.25] degrees, *θ* and *ϕ* from [0, *π*) and [0, 2*π*), respectively. Spatial frequency, *ω*, is drawn from a gamma distribution with shape parameter 2 and scale parameter 0.04 and 0.02 for excitatory and inhibitory neurons, respectively.

#### 1.2 Neuronal connectivity

Network connectivity is represented by the weight matrix *W*, with the entry *w*_*ij*_ denoting the weight of the connection from presynaptic neuron *j* to postsynaptic neuron *i*. Connectivity is all-to-all and weights between two neurons are modulated as a function of similarity of their respective receptive fields. Functional similarity is assayed in two ways: first, by calculating the correlation coefficient of the receptive fields (RF CC) directly:

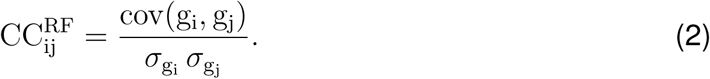

Alternatively, functional similarity is assayed by using external stimuli. Consider a sequence of stimuli, e.g. *N*_stim_ static gratings or natural images. The functional behaviour of neuron *i* in response to this stimulus set can be described by a response vector *ρ*_*i*_ (with size *N*_stim_). The *k*-th element of the vector represents the correlation coefficient of the neuronal RF with the *k*-th stimulus in the sequence. For a pair of neurons (*i, j*), the Response CC (also referred to as signal CC) is then calculated from the correlation of their response vectors, (*ρ*_*i*_, *ρ*_*j*_):

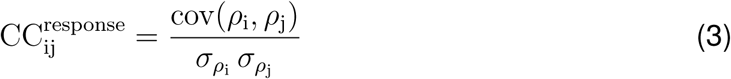

To evaluate response CCs in our simulations, we used 1000 static gratings with random preferred orientations between [0, *π*] and spatial frequencies drawn from a gamma distribution with shape parameter 2 and scale parameter 0.04. Gratings were instantiated in the same fashion as the Gabor RFs described above, with the difference that they were extended in space to obtain full-field gratings. This was achieved by choosing very large values of *σ* in Eq. (1).

For each pair of neurons *i* and *j*, the weight between them is then modulated as a function of the respective measure of similarity (CC_ij_):

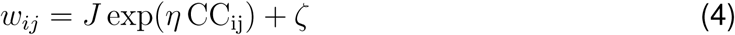

where CC_ij_ can be 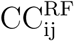 or 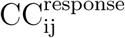 (this is specified in each case in the details of the simulation). J parameterises the strength of the respective weights (denoted as *J*_*XY*_, where {*X, Y*} ∈ {*E, I*}), and *η* determines the sharpness of the exponential dependence of weights on similarity (with a default value of *η* = 3). *ζ* is an i.i.d distributed randomly chosen value between [−*ζ*_*max*_, *ζ*_*max*_] added to each element, with *ζ*_*max*_ = 0.005. For *E* → {*E, I*} weights the values smaller than 0, and for *I* → {*E, I*} weights values larger than 0, are clipped to 0. For a given value of *E* → *E* weights (*J*_*EE*_), inhibition-dominance is parameterized by the relative surplus of *I* → {*E, I*} weights: *J*_*IE*_ = *J*_*II*_ = −*gJ*_*EE*_. The relative strength of *E* → *I* weights are quantified by a s imilarly d efined parameter, *α*: *J*_*EI*_ = *αJ*_*EE*_. Broadening of inhibitory connectivity (e.g. in **Figure 2D**) is controlled by changing the sharpness of the profile of *I* → {*E, I*} connections, by keeping *η* the same for other connections and changing *η*_*IE*_ and *η*_*II*_ specifically.

#### 1.3 Neuronal simulations

The activity of neuronal networks was simulated by numerically solving the following equations:

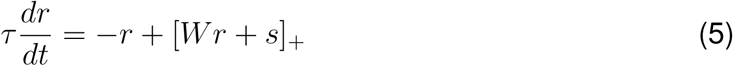

where *r* is a vector of all firing rates (for *N* neurons, composed of *N*_*E*_ excitatory and *N*_*I*_ inhibitory neurons), *W* is the matrix of connection weights as described above, *s* is the vector of external input to all neurons, *τ* is the effective time constant of integration, and []_+_ denotes half-wave rectification.

The influence is assayed by simulating the response of the network for a given time (*T*) in the baseline state (*s*_0_) and with an extra perturbation of a single neuron: *s*_*p*_ = *s*0 + *δs*_*i*_, where *δs*_*i*_ is a perturbation vector containing *δp* (the size of perturbation) for the i-th element and zeros for the rest of neurons. The average (temporal) firing rate of the neurons in the stationary state (after discarding the initial transient response for *T*_trans_) is calculated for the baseline and the perturbed state as *r*_0_ and *r*_*p*_, respectively. The vector of change in firing rates, *δr* = *r*_*p*_ − *r*_0_, is normalized by the size of perturbation to obtain the influence of neuron i (the influencer) on the rest of the network (influencees): *ψ* = *δr/δp*.

For double-neuron perturbations, the same procedure was repeated with the only difference that the vector of perturbations contained two non-zero elements for the two influencers with size *δp*, and influence was obtained by normalizing the induced changes in the activity of influencees by *δp*.

Default values for the quantification of influence are: *N*_*E*_ = *N*_*I*_ = 400, *τ* = 10, *T* = 500, *T*_trans_ = 50, *δp* = 0.1.

#### 1.4 Data analysis

To analyse the behaviour of the influence as a function of signal correlation between influencers and influencees in different regimes (**Figure 2**), we employed a feature-specific suppression/amplification (S/A) index. It was calculated from the average influence, which was obtained as the average influence between all pairs of influencers and influencees with signal correlations in a certain bin (with bin widths of 0.02) between −1 and 1. The index is composed of three submetrics: (1) *x*: the mean average influence in the intermediate regime; (2) *y*: the slope of the dependence of the average influence on signal correlation in the intermediate regime; and (3) *z*: the mean level of average influence in highly similar regime. The intermediate regime was defined as signal correlations between −0.3 to 0.3, and a highly similar regime was taken as the range of signal correlation between 0.7 to 0.9. Each submetric was normalized to the maximum of the absolute values for all the network simulations with different parameters tested (e.g. as in **Figures 2F-H**). The feature-specific S/A index (SAI) was then obtained as:

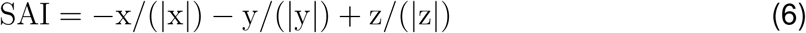

The more the suppression at intermediate regimes *and* the more the amplification at highly similar regimes, the higher the SAI would be.

To quantify the combined influence of perturbing two influencers in double-neuron perturbations (**Figure 4**), we developed a synergy index. For the first influencer (neuron *i*), the effect of additional perturbation of a second influencer (neuron *j*) on the influencee (neuron *k*) was quantified as follows:

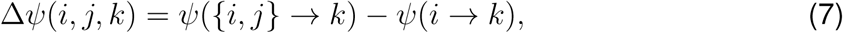

where *ψ*({*i, j*} → *k*) is the influence of double-neuron perturbations of {*i, j*} on *k* and *ψ*(*i* → *k*) is the single-neuron influence of *i* on *k*. The synergy of influence between the triplet {*i, j, k*} was then calculated as:

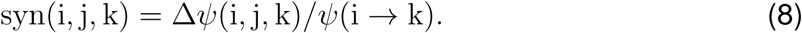

We excluded pairs with very small single-neuron perturbations (*ψ*(*i* → *k*) < 0.001) to avoid their overrepresentation in the metric. The average synergy between two influencers (*i, j*) (as in **Figures 4E,F**) was calculated by computing the mean synergy across all target influencees:

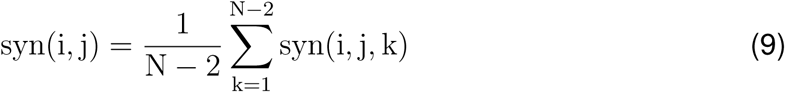

Note that the synergy will be positive (synergistic) if the change in the influence as a result of the interaction of the second influencer is in the same direction as the original, single-neuron perturbation, and negative (antagonistic) otherwise. Thus, both suppression and amplification of single-neuron influences can undergo synergy (or antagonism) as a result of double-neuron perturbations, depending on whether the interaction exacerbates (or diminishes) the initial influence in the same (or reverse) direction.

#### 1.5 Decoding of natural images

To evaluate the population responses of neuronal networks to external stimuli, we presented natural images to our model networks. Natural images were chosen from the McGill calibrated colour image database (http://tabby.vision.mcgill.ca). The feedforward input (*I*_ffw_) to each neuron in response to each natural image was calculated as:

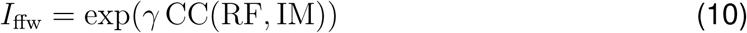

where CC(RF, IM) is the correlation coefficient of the image (IM) with the neuronal RF, and *γ* = 3 determines the sharpness of the exponential dependence. CC(RF, IM) was calculated for the central part of the image with the same size (in pixels) as RFs (that is, the central 200 × 200 pixels of the image for RFs instantiated on a visual field with 50 × 50 degrees extent and ppd = 4).

The activity of the network was calculated after accounting for recurrent operations on this input, by applying the matrix operator *A* = (*I* − *W*)^−1^ for different weight matrices:

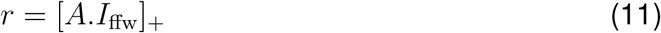

The gain of neuronal responses in response to each image is then obtained by dividing the activity of each neuron over its respective feedforward input. Discriminability of population responses to two images *i* and *j* are quantified by calculating the angle between the vectors of population responses (*r*_*i*_ and *r*_*j*_):

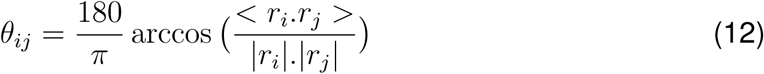

where <. > denotes the dot product and |.| is the norm of the vector. *θ*_*ij*_ = 90 corresponds to the maximum discriminability (orthogonal representations) and 0 or 180 degrees show the maximum collinear relationships (in the same or opposite directions, respectively). We use a normalized version of this angle:

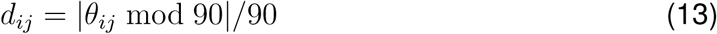

to quantify discriminability (as in **Figure 6H**), which ranges from 0 (minimum discriminability) to 1 (maximum discriminability).

To assess the decoding capacity of neuronal networks to discriminate natural images, we trained linear decoders on the population activity of the excitatory neurons. Each decoder was trained to distinguish a target image from other images. The ensemble of natural images was broken into two random parts (test and train sets, each containing 300 images) and the decoder was trained on the train set to detect the target image from the rest of the images. The training was done by presenting 300 pairs of images, containing the target image and one of the 300 test images. The decoder then finds (via linear regression) the best weighting of population activity of the network which separates the response to the target image from the non-target ones. To control for different levels of population activity under different conditions (e.g. networks with strong and weak recurrent interaction), we normalized the activity of the networks, such that the average activity of the network in response to each image was 1.

The decoder was tested on the test set, by presenting pairs of images containing the target image and each of the 300 test images. A threshold of 90% was set for the correct detection. The percent correct was then calculated as the fraction of the pairs for which the target image passed the threshold *and* the test image did not. The decoding task was performed for all 600 images as decoding targets. For each decoding task, the procedure was repeated for different levels of noise added to the population activity (both during training and for the test). It was added to the normalized activity of all excitatory neurons in response to each image and was drawn from a uniform distribution between 0 and *ξ*, with *ξ* ranging from 0.02 to 0.1 (**Figure 6K**). The example shown in **Figure 6J** had an intermediate noise level of 0.04.

### 2 Theoretical analysis of neuronal influence in single-neuron perturbations

We analytically evaluate the effect of single-neuron perturbations in networks of rate-based neurons as described above (Eq. (5)). We drop the firing threshold nonlinearity and analyze the linear behaviour of the network as described with the following dynamics:

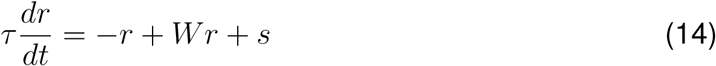

The stationary state solution of the firing rates under such dynamics is obtained under *dr/dt* = 0 and can be written as:

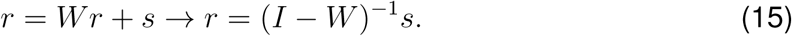

We define *A* = (*I* − *W*)^−1^ as the operator which acts on external input to obtain the steady-state firing rates in any equilibrium, *r*_0_ = *As*_0_. Single-cell perturbations around this steady-state leads to a new firing rate solution, *r*_*p*_ = *As*_*p*_. Here, *s*_*p*_ = *s*_0_ + *δs*, and *δs* is a vector of zeros at all entries except for the neuron which is perturbed. If the *i*-th neuron in the network is perturbed, we have:

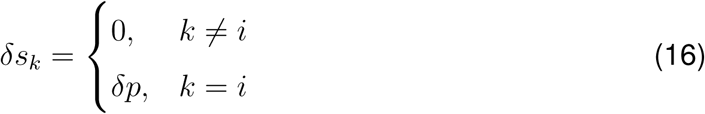

where *δp* is the size of perturbation. To obtain the influence of perturbation of neuron *i* on the postsynaptic neuron *j, ψ*(*i* → *j*) we need to calculate the change in the firing rate of the *j*-th entry of *r*_*p*_. Writing

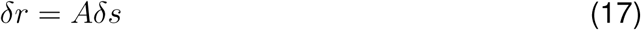

the rate change of the *j*-th neuron is obtained as:

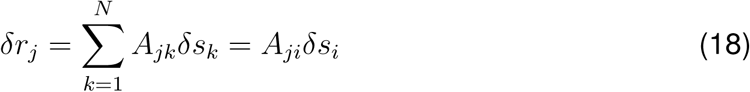

where *A*_*ji*_ is the entry on the *j*-th row and *i*-th column of matrix *A*. Writing the influence as the rate change of the influencee *j* divided by the perturbation strength of the influencer *i*:

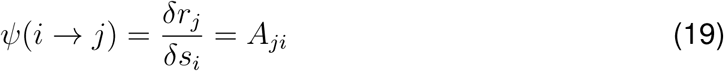

reveals that *A*_*ji*_ is, in fact, denoting the influence.

To obtain the neuronal influence in single-neuron perturbations, we used the above framework to evaluate *ψ*(*i* → *j*) = *A*_*ji*_, by mathematically calculating the influence in networks with different profiles of connectivity. We explain this approach in more detail below.

#### 2.1 Calculating influence for a general weight matrix

To obtain the influence, we calculate *A*_*ji*_, by expanding the matrix *A* with regard to *W* as:

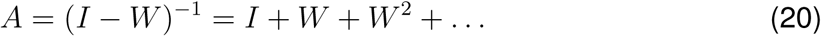

*A*_*ji*_ can therefore be expressed as:

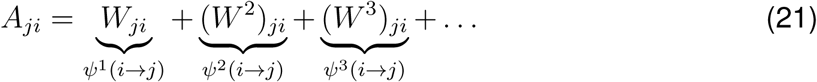

Note that the first term in Eq. (20) (the identity matrix) does not contribute to the influence in Eq. (21), since *i* ≠ *j* (the influencer and the influencee are different neurons).

The series describes different pathways of interaction from *i* to *j* in the following fashion:

I. Mono-synaptic influence denotes the direct interaction from *i* to *j*, which is inferred from the corresponding entry on the original weight matrix:

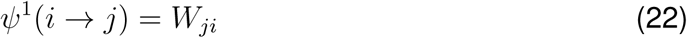
II. Di-synaptic influence entails second-order interactions, comprising all the pathways in which neuron *i* can influence neuron *j* via secondary neurons. It can be mathematically expressed as:

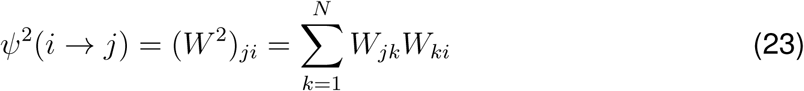

where index *k* denotes all the neurons in the network that mediate the influence from neuron *i* to neuron *j*.
III. Tri-synaptic influence captures all interactions with two layers of intermediate neurons, denoted by indices *k* and *l* in the following formulation:

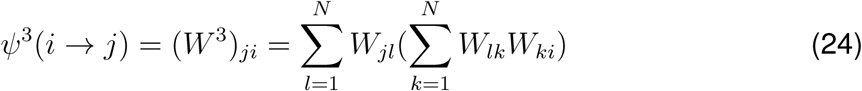

Higher order interactions (including tetra-synaptic *ψ*^4^(*i* → *j*), penta-synaptic *ψ*^5^(*i* → *j*), etc) can be calculated via similar equations.

##### 2.1.1 Networks with excitation and inhibition

In the next step, we calculated the influence for networks containing two subtypes of excitatory (*E*) and inhibitory (*I*) neurons, with the number of neurons in the network denoted by *N*_*E*_ and *N*_*I*_, respectively. The connection weights from E to E, E to I, I to E and I to I neurons are described, respectively, by *J*_*EE*_, *J*_*EI*_, *J*_*IE*_, and *J*_*II*_. *W* is ordered such that the first *N*_*E*_ elements are excitatory neurons and the next *N*_*I*_ elements (*N*_*E*_ +1 to *N*_*E*_ +*N*_*I*_ rows/columns) represent inhibitory neurons.

The influence of the *i*-th excitatory neuron on the *j*-th excitatory neuron in the network can be calculated as:

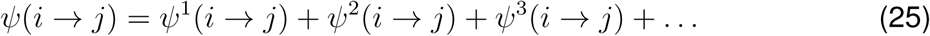

where different orders of influence are calculated as the following:

I. Mono-synaptic influence is given by

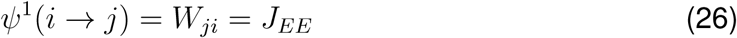

which is the direct connection between the two excitatory neurons.
II. Di-synaptic influence is calculated as

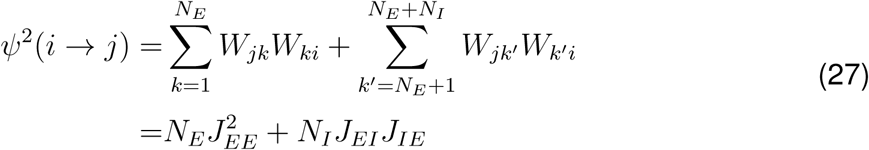

which contains pathways with either excitatory or inhibitory neurons in between the influencer and the influencee.
III. Tri-synaptic influence can, in turn, be written as

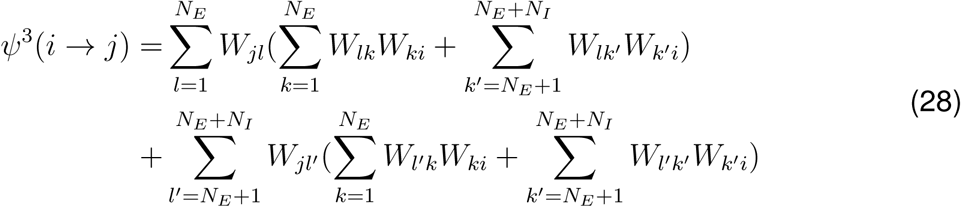 This includes four possibility of mediation between two excitatory neurons, *E* → *X* → *Y* → *E* (namely *E* → *E* → *E* → *E, E* → *E* → *I* → *E, E* → *I* → *E* → *E, E* → *I* → *I* → *E*, respectively) and can be calculated as:

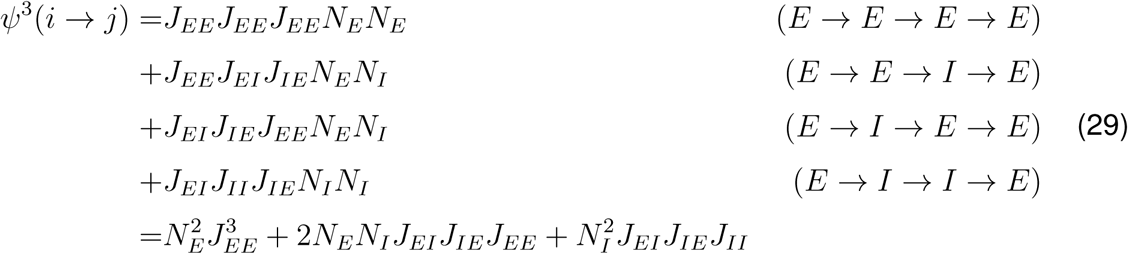 Note that the latter motif entails a net positive influence, as it involves inhibition of inhibition.
IV. Tetra-synaptic influence can, similarly, be mediated by 3-order motifs (*E* → *X* → *Y* → *Z* → *E*, where {*X, Y, Z*} can be either *E* or *I*, leading to a total of 8 possibilities), and therefore can be written as

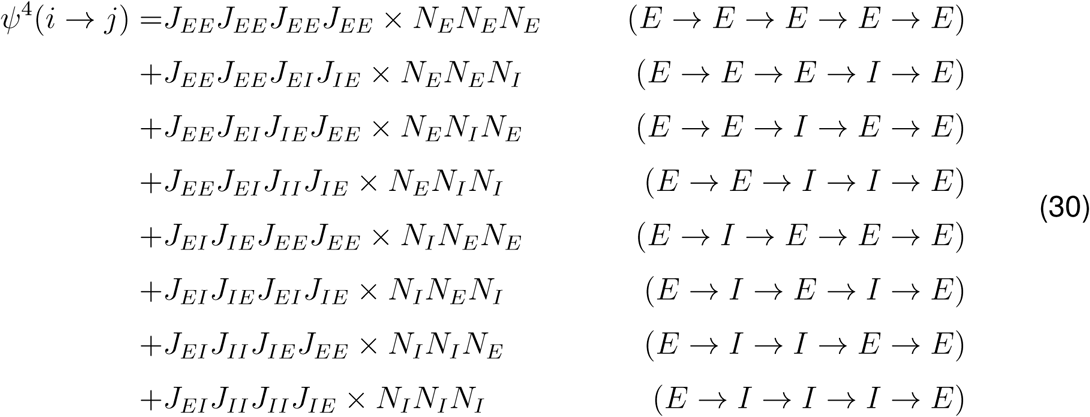

Higher order influences can be calculated in a similar fashion by counting higher-order motifs.

##### 2.1.2 The case of inhibition-dominance

It is useful to calculate the influence for a simplified description of the abovementioned weight matrices, where *J*_*EE*_ = *J, J*_*EI*_ = *αJ*_*EE*_, and *J*_*IE*_ = *J*_*II*_ = −*gJ*_*EE*_. We further assume *N*_*E*_ = *N*_*I*_ = *N* and write:

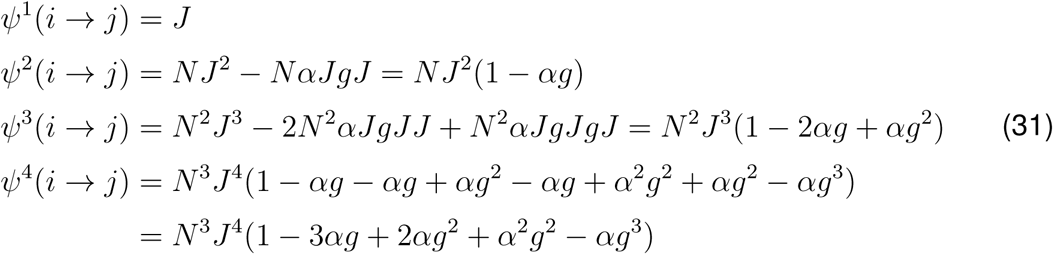

For the specific condition that *α* = 1, we have:

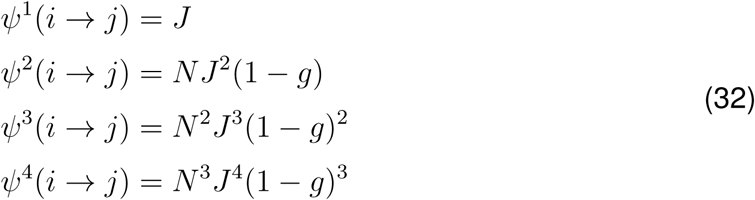

which in fact provides a closed-form description of the influence at all orders of influence:

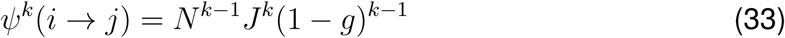

The total influence can therefore be written as:

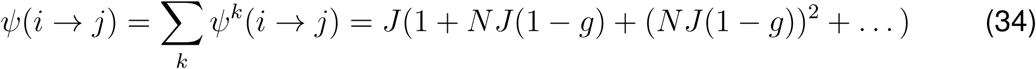

which can be expressed as:

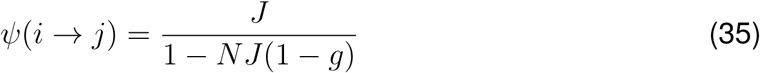

Note that inhibition dominance, *g* > 1, does *not* imply a negative influence here. Rewriting *κ* = 1 − *NJ* (1 − *g*), and noticing that *κ* > 1 for *g* > 1, we observe *divisive inhibition* as a result of inhibition dominance:

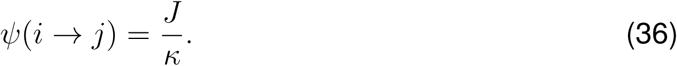

The stronger the inhibition-dominance, the larger the divisive term in the denominator, and hence the higher the divisive inhibition.

#### 2.2 Calculating influence for networks with specific connectivity

The calculations presented in the previous section can be extended to networks with specific connectivity, where the weight of connections between neurons are defined as a function of their functional similarity. First we consider a scenario where the functional property of neurons is defined by a one-dimensional parameter, e.g. their preferred orientations, *θ*. We consider weight matrices described by:

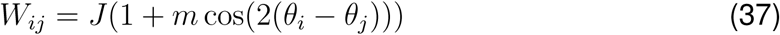

where the connection weight between neuron *i* and *j* is modulated by the similarity of their respective preferred orientations. *m* determines the degree of specificity of connections, with *m* = 0 retrieving the unspecific weight matrices described in the previous section. We now calculate the influence (*A*_*ji*_) for a network of excitatory and inhibitory neurons with specific connectivity, described as:

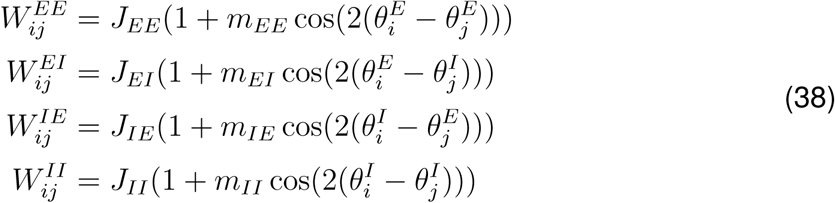

Here, *J*_*XY*_ and *m*_*XY*_ denote, respectively, the average weight and the degree of specificity of synapses, and *θ*^*X*^ represents the preferred orientation. (*X, Y*) ∈ (*E, I*).

We first calculate the influence for a scenario where all connections have the same degree of specificity, i.e. *m* _*EE*_ = *m* _*EI*_ = *m* _*IE*_ = *m* _*II*_ = *m*. We also assume that *J* _*EE*_ = *J, J*_*EI*_ = *αJ*_*EE*_, and *J*_*IE*_ = *J*_*II*_ = −*gJ*_*EE*_, as described above. Under these conditions, the influence of perturbing excitatory neuron *i* on the excitatory neuron *j* can be calculated as the following for different orders of interaction:

I. Monosynaptic:

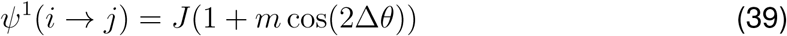

where Δ*θ* = *θ*_*i*_ − *θ*_*j*_.
II. Di-synaptic:

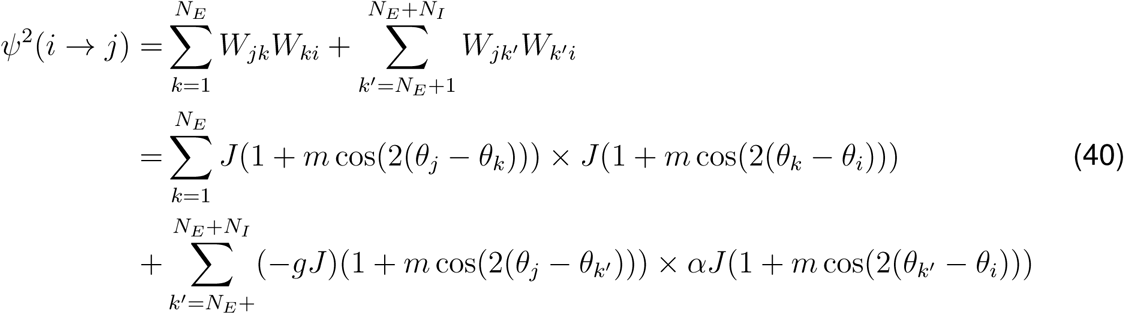 Assuming large *N*, we can solve the following continuous version of the equation:

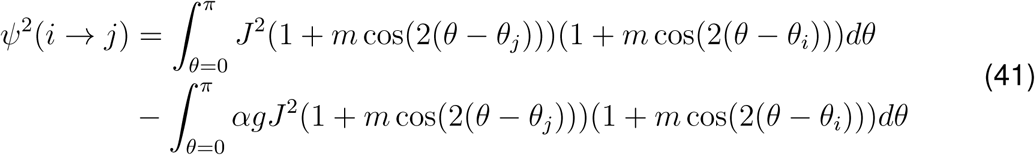

which, given Δ*θ* = *θ*_*i*_ − *θ*_*j*_, leads to

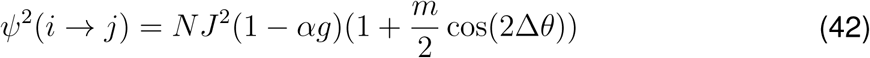 In calculating the above identity, we used the following equation:

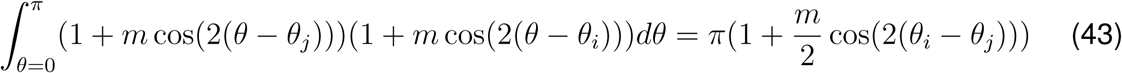

and its discrete equivalent:

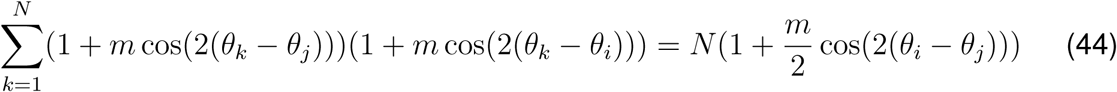 It describes how indirect weights and their specificity are effectively determined when a layer of intermediate neurons is mediating the influence, and is useful to highlight since it appears recurrently in calculating higher-order motifs in what follows.
III. Tri-synaptic:

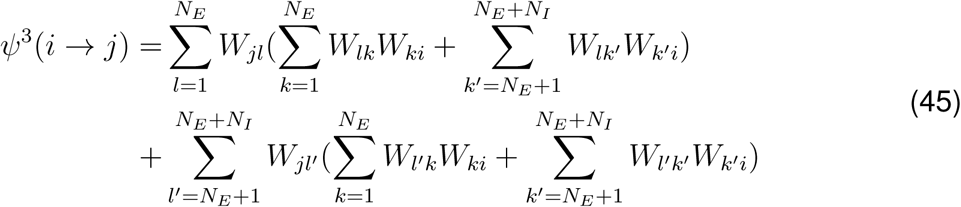

which can be written as:

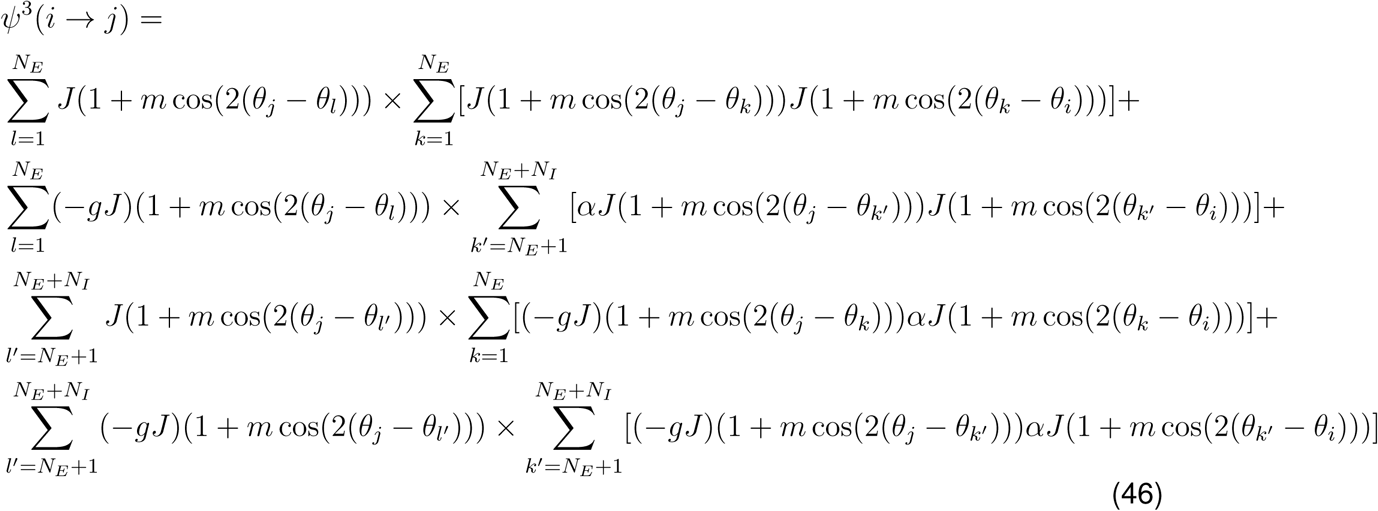

and results in:

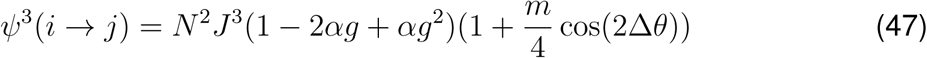
IV. Tetra-synaptic: Accounting for all fourth order motifs in a similar fashion as explained above, we obtain the following for the fourth order influence:

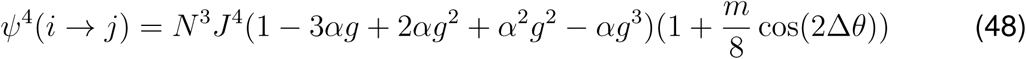 The total influence between excitatory neurons *i* and *j* in specific EI networks can, therefore, be expressed as:

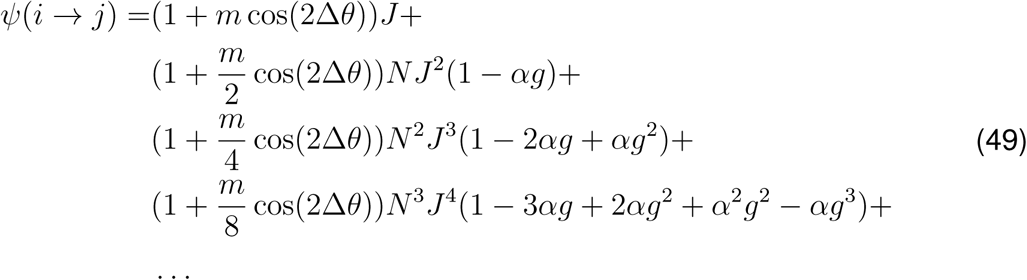 Note that the influence contains nonspecific and specific components, and that the nonspecific component is similar to what we obtained before for influence in nonspecific networks (Eq. (31). For the simplified case of *α* = 1, we can follow similar steps as described above for nonspecific networks to characterize the influence for all higher orders with a closed form expression:

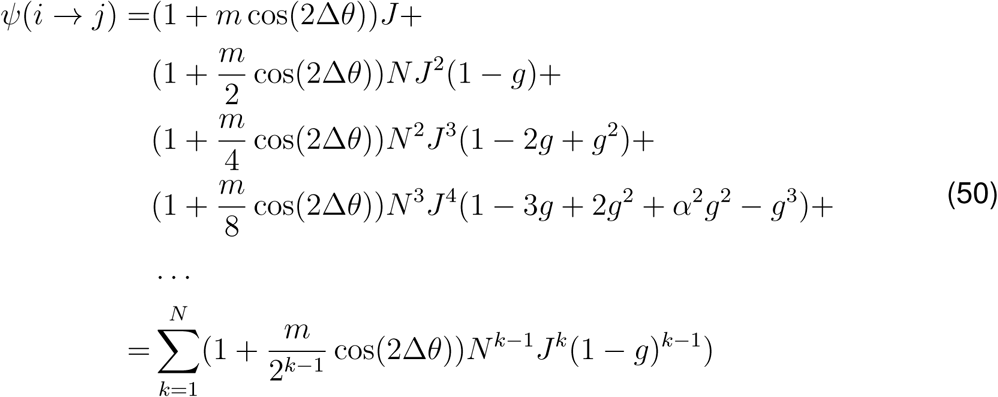 The nonspecific part retrieves the same formulation as before (c.f. Eq. (35)):

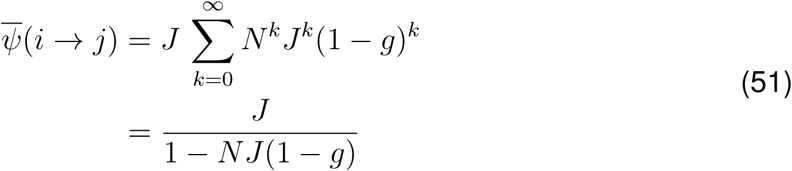

while the specific component of the influence can be expressed as:

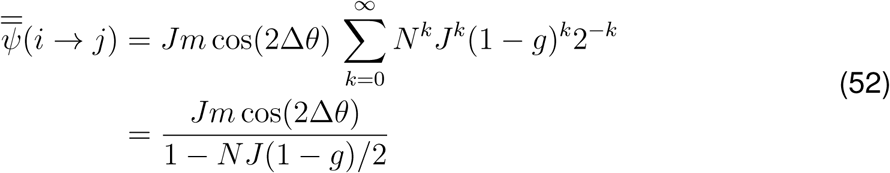 Note that inhibition dominance (*g* > 1) does *not* yield a net negative influence here too. That is, *feature-specific suppression* does not result from general inhibition dominance in specific EI networks, when *α* = 1. Instead, it leads to (feature-specific) divisive inhibition (c.f. Eq. (35)), with higher inhibition-dominance values (*g*) implying higher divisive terms for the specific component.

##### 2.2.1 Networks with broad inhibitory connectivity

So far, we considered the scenario where all connections had the same degrees of specific connectivity, by assuming *m*_*EE*_ = *m*_*EI*_ = *m*_*IE*_ = *m*_*II*_ = *m*. We now relax that assumption by allowing excitatory and inhibitory weights to have different degrees of specificity. We assume *m*_*EE*_ = *m*_*EI*_ = *m*_*e*_ and *m*_*IE*_ = *m*_*II*_ = *m*_*i*_, and solve for the condition where inhibition has a broader (i.e. less specific) connectivity, *m*_*e*_ > *m*_*i*_.

Accounting for broader inhibition does not change the nonspecific component of the influence, but the specific component can now be written as:

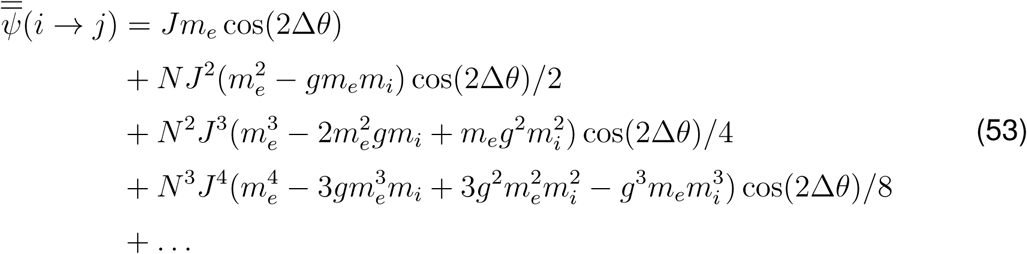

Defining *J*′ = *m*_*e*_*J* and *g*′ = *gm*_*i*_*/m*_*e*_, we can write:

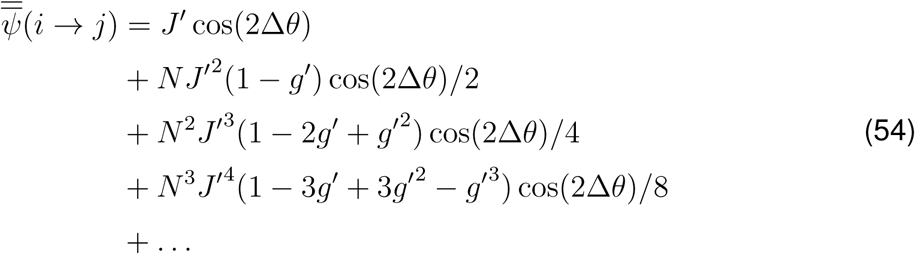

and therefore the specific component of the influence can be calculated as

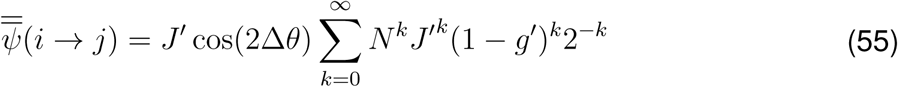

which leads to:

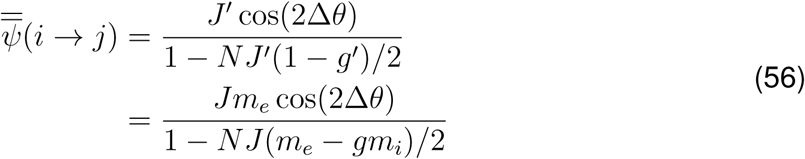

To obtain feature-specific suppression in single-neuron influences (namely, more suppression for pairs with smaller Δ*θ*), we need to have 1 − *NJ* (*m*_*e*_ − *gm*_*i*_)*/*2 < 0. Broader inhibition does not confer such a negative influence in inhibition-dominance networks, if *g m*_*i*_ > *m*_*e*_. The only situation under which such a negative influence appears is if *m*_*e*_ > *gm*_*i*_ and *NJ* (*m*_*e*_ − *gm*_*i*_)*/*2 > 1 at the same time, but note that the latter condition implies instability of the weight matrix along the specific eigenmode.

#### 2.3 Calculating influence for specific E-I networks with strong E-to-I connectivity

In this section, we relax the previously made assumption of *J*_*EE*_ = *J*_*EI*_, and allow the excitatory neurons to have different connection weights to their excitatory and inhibitory postsynaptic targets, formulated by: *J*_*EE*_ = *J, J*_*EI*_ = *αJ*_*EE*_, *J*_*IE*_ = *J*_*II*_ = −*gJ*_*EE*_. We assume similar connection specificity for all synapses: *m*_*EE*_ = *m*_*EI*_ = *m*_*IE*_ = *m*_*II*_ = *m*. Following similar procedures as described before for networks with specific connectivity (and summarized in Eq. (49)), different orders of influence in specific networks with strong *E* → *I* connectivity can be written as:

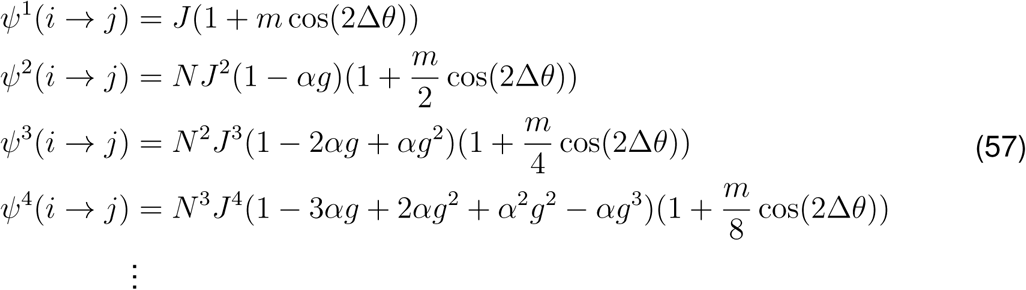

We define *ψ*_*E*_ = *ψ*_*E→X→E*_ and *ψ*_*I*_ = *ψ*_*I→X→E*_ as two second-order, di-synaptic motifs of influence, one starting from an excitatory and the other from an inhibitory neuron, respectively. The excitatory second-order motif *ψ*_*E*_ is *ψ*^2^(*i* → *j*), by definition, and can hence be written as before:

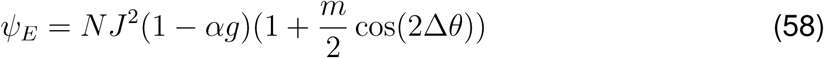

The inhibitory second-order motif can, in turn, be calculated as:

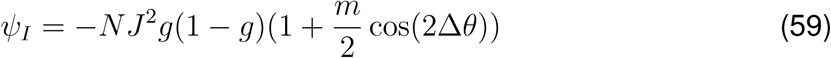

Note that, although starting from an inhibitory neuron, this motif would be positive for *g* > 1 (the condition which we referred to as inhibition-dominance). That is, inhibition-dominance leads to a net *excitatory* effect of the second-order motif of inhibitory neurons; this is because the net positive effect of inhibition of inhibition (*I* → *I* → *E*) is larger than the net negative effect of inhibition of excitation (*I* → *E* → *E*).

We define

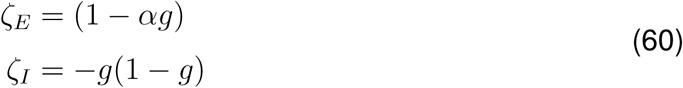

as the main factors appearing in the second-order excitatory and inhibitory motifs. This allows us to write the specific component of the higher-order motifs (in Eq. (57)) in terms of the basic factors of the respective di-synaptic motifs:

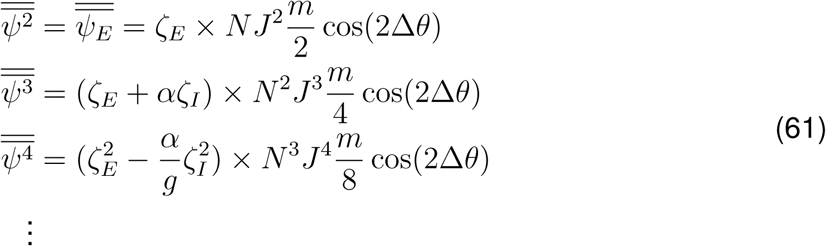

Feature-specific suppression implicates a higher suppressive influence between neuronal pairs with smaller Δ*θ*. We can now evaluate this for specific components of higher-order motifs of influence, in view of the interaction of the basic di-synaptic motifs.

[1] For the 2nd-order motif, this implies that the basic excitatory second-order motif is negative, hence:

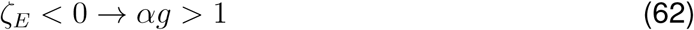

Both *E* → *I* connections (parameterized by *α*) and *I* → {*E, I*} weights (parameterized by *g*, or inhibition-dominance) can be strong to satisfy this condition, which highlights the significance of the specific excitatory-inhibitory interaction.

[2] For networks with weak recurrent coupling (*NJ* ≪ 1), higher-orders of influence would be much smaller than the lower orders and hence can be ignored in the net influence. However, higher-order motifs cannot be ignored in networks with strong coupling (*NJ* ≫ 1). In that case, the condition inferred from the 2nd-order motif, *ζ*_*E*_ < 0, does not guarantee a negative specific 3rd-order motif, since we need: (*ζ*_*E*_ + *αζ*_*I*_) < 0. However, as we argued above, inhibition-dominance (*g* > 1) implies a positive *ζ*_*I*_. *ζ*_*I*_ is only negative if we have:

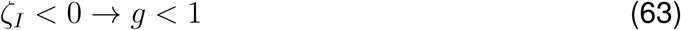

This means that *α* and *g* cannot be arbitrarily increased as we assumed for the 2nd-order motif in [1].

An alternative way to satisfy this condition might be obtained by broadening of inhibition. If we allowed for different selectivity of excitatory and inhibitory connectivity (as denoted by *m*_*e*_ and *m*_*i*_ in Section. 2.2.1), the condition in Eq. (63) would change to *g m*_*i*_*/m*_*e*_ < 1. Now, this can be satisfied by changing *g* and/or *m*_*i*_*/m*_*e*_, with the small values of the latter (*m*_*i*_*/m*_*e*_ < 1) implying broaderer specificity of inhibitory connections compared to excitation. Overall, it means a weak specific inhibitory connectivity, which can be satisfied by weaker or broader inhibition. As strong inhibition-dominance might be necessary to balance nonspecific excitation, broader inhibition might be a better strategy to satisfy the negative influence of this specific motif.

[3] So far, negative influence of motifs could be satisfied, if the the two basic 2nd-order excitatory and inhibitory specific motifs were negative. For the 4th-order specific motif to be negative, we need:

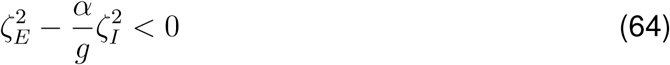

We therefore have:

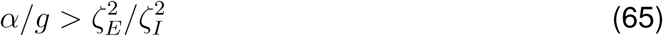

One way to achieve this is to have balanced di-synaptic motifs: *ζ*_*E*_ ≈ *ζ*_*I*_. Under this condition, we have 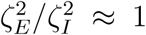, and we therefore need *α/g* > 1, or *α* > *g*. But this is already satisfied, since the two conditions *αg* > 1 (from Eq. (62)) and *g* < 1 (from Eq. (63)), imply *α* > 1 > *g* and hence *α* > *g*.

Taken together, several conclusions can be inferred from our little exercise here: first, it is not possible to obtain negative influence for specific motifs by simply increasing the inhibition-dominance *g*; too large values of *g* can lead to strong disinhibitory effects, which might counteract the direct inhibitory effects via higher-order interactions. Second, the strength of *E* → *I* connections are as important, if not more, to achieve negative influence. Finally, balancing excitatory and inhibitory motifs might be needed, to avoid the dominance of one over the other.

However, although illustrative, these results are not conclusive, for the following reasons. First, we do not have a closed-form expression for higher-order motifs, which hinders a systematic evaluation of all such terms and the conditions for their negativity. Moreover, the actual condition for the net negative influence is not the negativity of each higher-order term, but the negativity of the net sum. We therefore need mathematical formulations which provide us with these two requirements. We attempt to provide such analyses in the following sections.

##### 2.3.1 Networks with dominant E-to-I connections only

It is instructive to study a special case where *α* ≫ 1 and *g* = 1, to evaluate the effect of strong E-to-I connections in the absence of strong inhibition-dominance. Under this condition, the basic di-synaptic motifs (Eq. (60)) can be written as

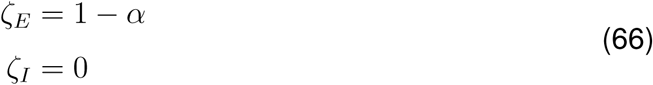

Only the excitatory component of basic di-synaptic motifs counts for calculating higher-order motifs, and we can therefore express different motifs as the following:

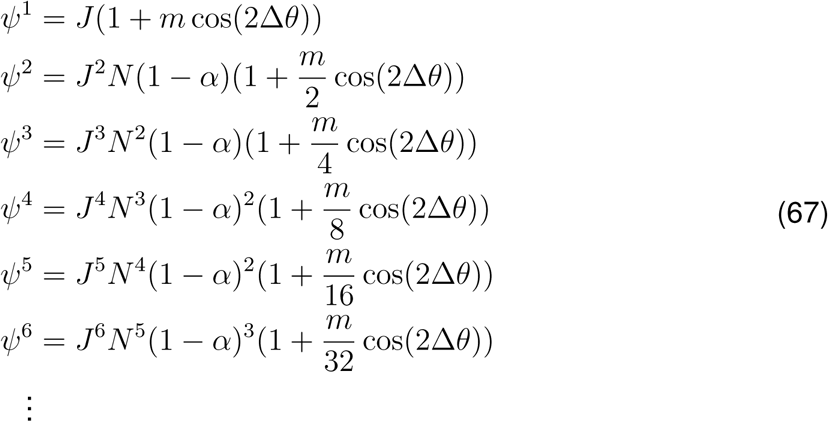

Writing in terms of the sum of subsequent motifs (*ψ*_1_ + *ψ*_2_, *ψ*_3_ + *ψ*_4_, *ψ*_5_ + *ψ*_6_, …), we reach to the following closed-form expression of the total influence, for nonspecific and specific components, respectively:

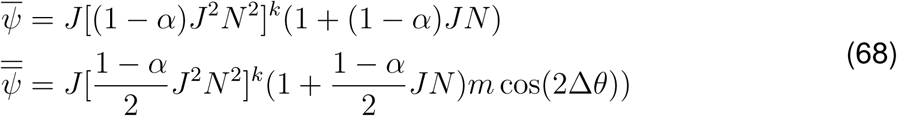

and hence:

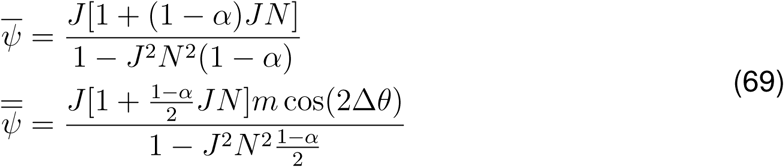

If *α* > 1, the denominator is positive, and the condition for a net negative influence becomes:

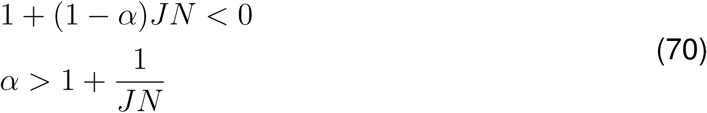

for the nonspecific, and

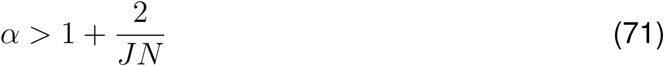

for the specific component of the influence. Both conditions can be met by strong E-to-I (*α*) and/or strong connectivity (*JN*), but *α* should be stronger for a net negative specific influence.

##### 2.3.2 Solution for strong E-to-I and inhibitory connections

Here, we consider a more general conditions, where *E* → *I* and *I* → {*E, I*} connections are both arbitrary and strong (i.e. *J*_*EE*_ = *J, J*_*EI*_ = *αJ*_*EE*_, *J*_*IE*_ = *J*_*II*_ = −*gJ*_*EE*_, and *g* > 0, *α* > 0). We start by writing the *n*-th excitatory and inhibitory motifs of influence (i.e., the motifs starting from an *n*-th order *E* or *I* neurons in the chain of influence), in terms of the subsequent motifs:

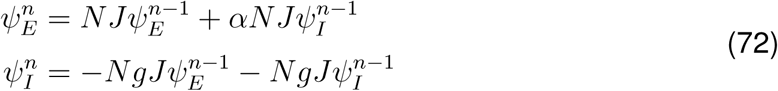

Replacing inhibitory motifs from the second equation iteratively, the *n*-th excitatory motif can be written in terms of lower order excitatory motifs as:

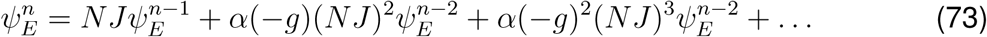

which can be described by the following recursive formula:

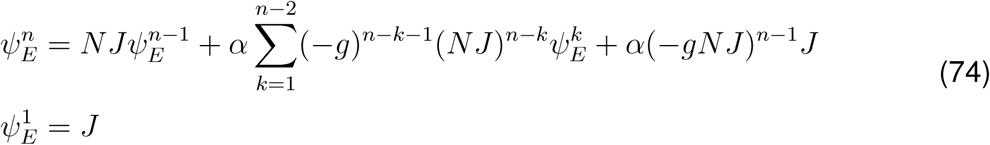

If we write Eq. (72) in the matrix form:

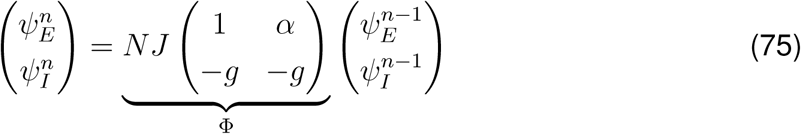

the *k*-th motif can be obtained by applying the matrix Φ, recursively, on lower orders:

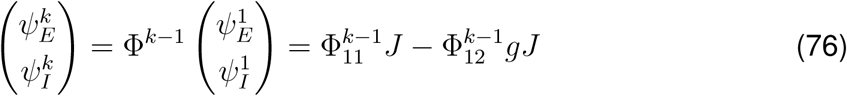

Here, 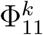 and 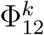 are the entries on the first row and the first and the second columns, respectively, of the *k*-th power of Φ, and we have used the identities 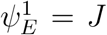 and 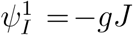. The influence of an excitatory neuron *i* on a target neuron *j* can now be calculated by counting the influence via all such higher order motifs:

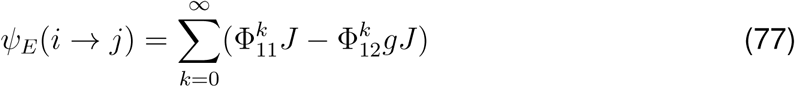

Defining 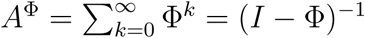, we can write:

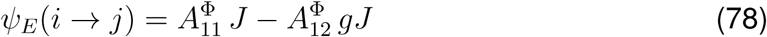

*A*^Φ^ can be calculated as:

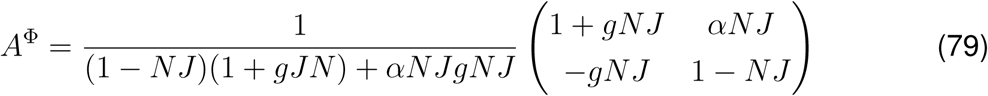

and hence the influence can be written as:

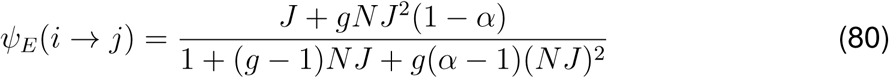

Strong *g* and *α* would make the denominator positive, but for a negative influence the numerator should be negative, which, assuming strong connectivity and hence *gNJ* ≫ 1, necessitates the condition of *α* > 1. Hence, strong *E* → *I* connections is necessary to obtain a negative influence. Similar arguments can be made for the tuned component of influence, which argues for the necessity of strong *E* → *I* specific connectivity in order to obtain feature-specific suppression in single-neuron perturbations.

#### 2.4 Solution for general connectivity conditions

Here, we consider the most general condition, where all connections (*E* → {*E, I*} and *I* → {*E, I*}) can have arbitrary weights. We parameterize this condition by defining: *J*_*EE*_ = *J, J*_*EI*_ = *αJ*_*EE*_, *J*_*IE*_ = −*βgJ*_*EE*_, and *J*_*II*_ = −*gJ*_*EE*_. Dominance of weights with regards to the excitatory weight, *J*_*EE*_, is thus determined by three independent factors: *α* denoting the dominance of *E* → *I*; *g* denoting the dominance of inhibition; and *β* parameterizing the extra inhibition of *I* → *E* weights. All factors can indicate dominance (> 1) or lack thereof (< 1).

The *n*-th excitatory and inhibitory motif of influence can be expressed now in terms of previous motifs (similar to Eq. (72)), in a recursive fashion:

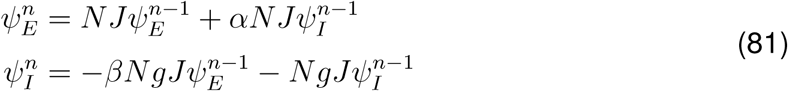

Expressing in matrix form as before (Eq. (75)), we have:

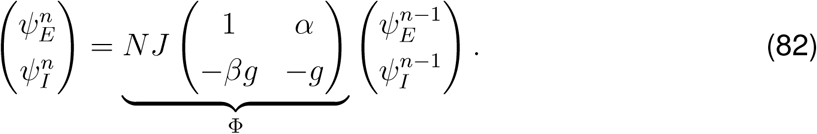

The *k*-th order motif can now be obtained by recursive application of operator Φ on all lower-order (< *k*) motifs, resulting in:

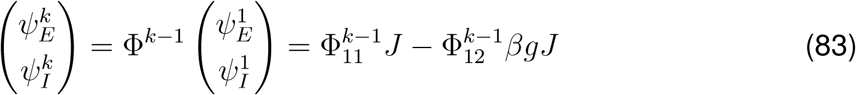

The influence of an excitatory neuron *i* on a target neuron *j* can again be calculated by counting the influence via all higher order motifs:

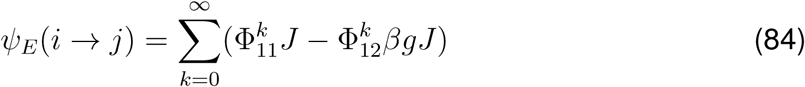

which can, in turn, be written as:

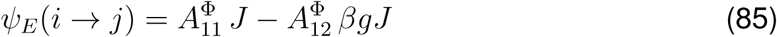

where

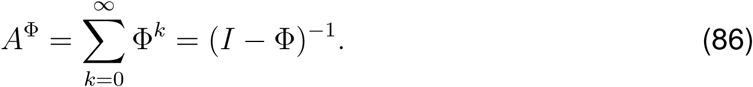

*A*^Φ^ can now be calculated as:

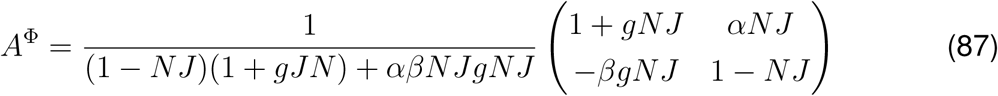

The influence can therefore be expressed as:

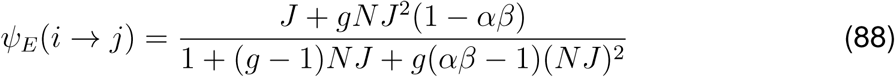

Following the same argument made for Eq. (80), we reach to the conclusion that, given a strong level of inhibition-dominance (*g* ≫ 1), the condition for negative influence is *αβ* > 1. This argues for a strong interaction of dominant *E* → *I* and *I* → *E* weights as the necessary condition for suppressive influence of single-neuron perturbations. Similar formulation and argument for the case of specific connectivity argues for strong and specific connectivity of *E* → *I* and *I* → *E* as a prerequisite for suppressive influence along the specific dimension, and hence *feature-specific suppression*. This, in turn, explains why global inhibition dominance, or broad inhibition, alone is not enough for feature-specific suppression in EI networks.

